# A comprehensive multi-omics study reveals potential prognostic and diagnostic biomarkers for colorectal cancer

**DOI:** 10.1101/2024.06.10.598127

**Authors:** Mohita Mahajan, Subodh Dhabalia, Tirtharaj Dash, Angshuman Sarkar, Sukanta Mondal

## Abstract

**Background:** Colorectal cancer (CRC) is a complex disease with diverse genetic alterations and causes 10% of cancer-related deaths worldwide. Understanding its molecular mechanisms is essential for identifying potential biomarkers and therapeutic targets for its effective management.

**Method:** We integrated copy number alterations (CNA) and mutation data via their differentially expressed genes termed as candidate genes (CGs) computed using bioinformatics approaches. Then, using the CGs, we perform Weighted correlation network analysis (WGCNA) and utilise several hazard models such as Univariate Cox, Least Absolute Shrinkage and Selection Operator (LASSO) Cox and multivariate Cox to identify the key genes involved in CRC progression. We used different machine-learning models to demonstrate the discriminative power of selected hub genes among normal and CRC (early and late-stage) samples.

**Results:** The integration of CNA with mRNA expression identified over 3000 CGs, including CRC-specific driver genes like *MYC* and *APC*. In addition, pathway analysis revealed that the CGs are mainly enriched in endocytosis, cell cycle, wnt signalling and mTOR signalling pathways. Hazard models identified four key genes, *CASP2, HCN4, LRRC69* and *SRD5A1*, that were significantly associated with CRC progression and predicted the 1-year, 3-years, and 5-years survival times. WGCNA identified seven hub genes: *DSCC1, ETV4, KIAA1549, NOP56, RRS1, TEAD4* and *ANKRD13B*, which exhibited strong predictive performance in distinguishing normal from CRC (early and late-stage) samples.

**Conclusions:** Integrating regulatory information with gene expression improved early versus late-stage prediction. The identified potential prognostic and diagnostic biomarkers in this study may guide us in developing effective therapeutic strategies for CRC management.

## 1. Introduction

Global Cancer Observatory statistics suggest that colorectal cancer (CRC) is a leading cause of cancer-related death worldwide, ranking third among all cancer types [1] and accounts for approximately 10% of all cancer-related deaths worldwide [2]. Some potential reasons explaining this high incidence could be an ageing population and unhealthy lifestyles, such as poor dietary habits, smoking, low physical activity [3]. Over the last decade, CRC has been one of the most growing field for healthcare studies and attracting scientific attention from all areas of science, including artificial intelligence [4, 5]. In all studies, there has been a consensus on one fact that CRC is a complex and heterogeneous disease involving a diverse range of genetic alterations and abnormal gene expression. For example, approximately 90% of CRC cases exhibit genetic alterations in the *APC* gene, a key driver gene involved in the initiation and progression of CRC [6, 7]. While 85% of the CRC cases exhibit a mutation in tumour suppressor genes such as *KRAS, PIK3CA*, and *TP53* [8]. These genomic changes also affect gene expression and the biological pathways and help in the development of tumours [9]. We need to understand the underlying molecular mechanism at the systems level to identify effective biomarkers and therapeutic targets.

Advanced high-throughput technologies provide molecular data from different levels of central dogma, such as genomics, transcriptomics, and proteomics. Analysing these “omics” data can help to understand the underlying molecular mechanism of complex diseases such as cancer [10]. Integrating data from different omics levels, often collectively referred to as “multi-omics” can provide excellent opportunities to understand disease mechanisms at the system level. In recent years, many studies have integrated different omics data to gain insights into the underlying mechanisms and identify potential therapeutic targets for diseases. For example, Liu et al. in 2021 integrated copy number alterations with transcriptomic data to identify key driver genes in gastric cancer [11] and showed their significant association with the clinical outcome, such as overall survival in gastric cancer. Similarly, Shao et al. integrated genetic alterations with mRNA expression to understand their relationship in pan-cancer studies [12]. Using integrative multi-omics analysis, Sivadas et al. [13] identified a molecular signature to distinguish invasive lobular carcinoma from invasive breast cancer. [14] used a multi-omics approach to classify the CRC into two distinct subtypes with different prognoses. They also identified the association of the MID2 gene with the progression of CRC stages, showing its impact on epithelialmesenchymal transition and the aggressiveness of CRC cells through experimental validation.

The continuing success of multi-omics studies on complex diseases like cancer motivated our research in this paper to investigate its effectiveness in providing some novel understanding of colorectal cancer (CRC) at the molecular level. Specifically, we conduct a comprehensive computational investigation of CRC using multi-omics data integration, with an aim towards discovering biomarkers for the prognosis and diagnosis of CRC. The major contributions of our study are outlined below.

1. We integrate genomic information, including copy number variation and somatic mutation, with the gene expression and clinical information of samples to understand the initiation and progression of CRC.
2. We compute differentially expressed genes to construct a set of candidate genes (CGs) that may play a crucial role in CRC progression. We then perform functional enrichment analysis to understand the biological significance of these identified CGs in the context of CRC. Furthermore, we use several hazard models to compute a minimal subset of key driver genes involved in CRC progression.
3. We explore the co-expression relationships of the CGs using Weighted Gene Co-expression Network Analysis (WGCNA) to identify the key hub genes that show significant associations with clinical traits, such as early and late stages of CRC. Here, we also use several machine learning models, both unsupervised and supervised, to demonstrate the discriminative power of these hub genes in predicting disease status.

The rest of the paper is organised as follows: In section 2, we start with a description of our data collection steps, followed by various methodologies of our computational models, such as WGCNA, gene regulatory network and machine learning. We provide all our key results in section 3 and discuss our findings broadly in section 4. We conclude our paper in section 5. We then provide public links to how to use our data and codes for further potential use cases. We share all our codes and data via GitHub.

## 2. Methods

The overall workflow used in our study is presented in Figure 1 that shows several stages of our multi-omics analysis pipeline. Each of these stages is elaborated in the following sections.

**Figure 1:**
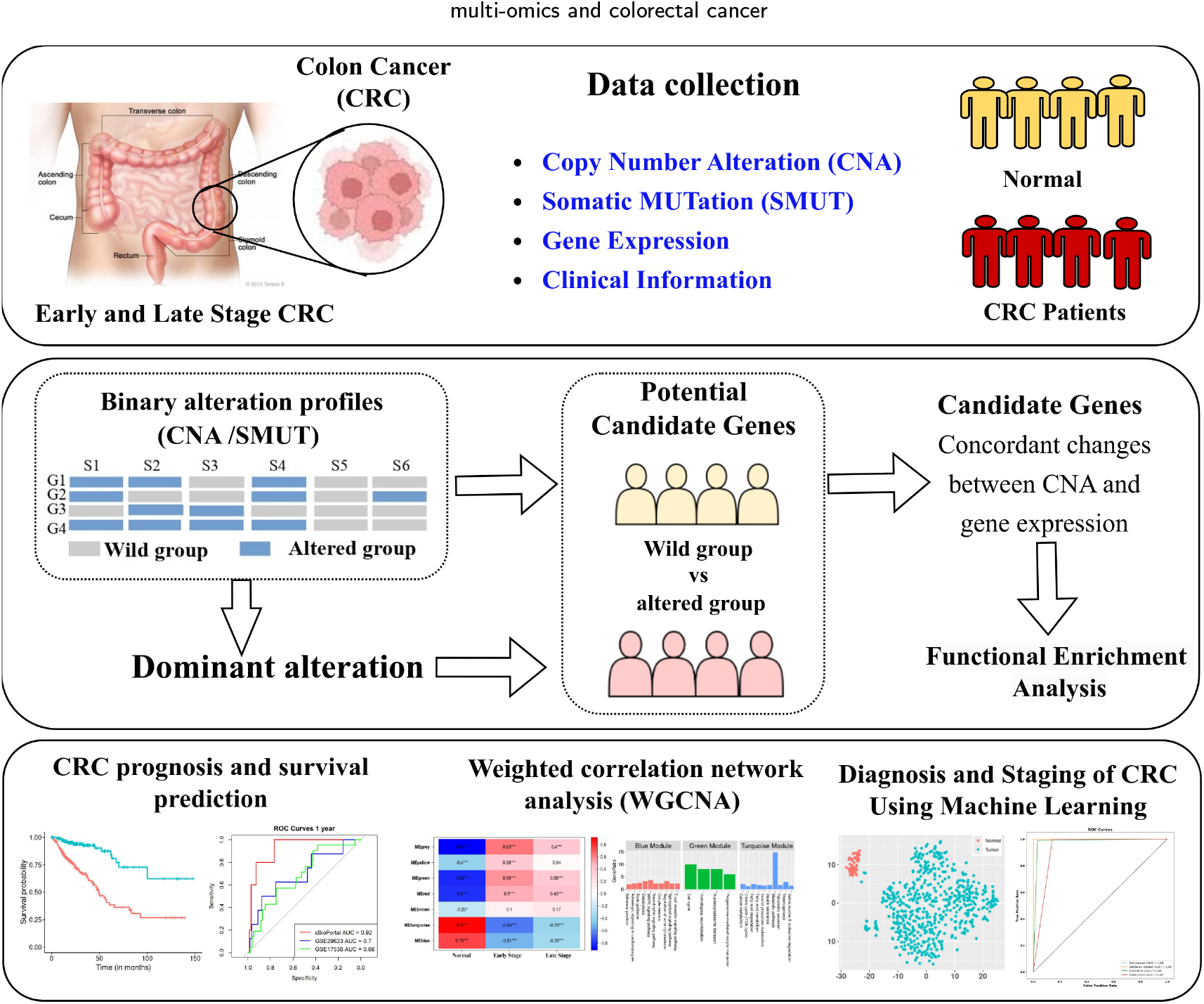
An overview of this study. The first stage is to collect relevant datasets across different layers of central dogma: Genomics, transcriptomics and clinical. The second stage consists of bioinformatics analyses, including differential expression to construct a set of candidate genes (CGs). The next stage involves a more complex analysis of these CGs: prognostic analyses, gene regulatory and pathway analyses and diagnostic analyses using machine learning.

## 2.1 Data collection and processing

The multi-omics data, including CNA, mutation data and gene expression (RNA-Seq V2 RSEM), were collected from The Cancer Genome Atlas (TCGA) and Firehose Legacy through cBioPortal data sources [15, 16]). A summary of information on the multi-omics data used in our study is provided in Table 1. Genomic Identification of Significant Targets in Cancer (GISTIC 2.0 [17]) was used to determine the copy number alterations, and it was then discretised into five types: homozygous deletion (−2), heterozygous deletion (−1), no change (0), gain (1) and amplification (2) for each gene in all cancer samples. The numbers in brackets represent the copy-number level per gene. The gene expression datasets used were normalised RSEM (RNASeq by Expectation-Maximization) values from the RNAseq V2, and we performed log transformation (that is, *x* ↦ log_2_(*x* + 1)) for the downstream analysis, where *x* is the normalised RSEM value. For mutation data, we considered the somatic mutation to have a valid status and excluded silent mutation. For mRNA, the genes expressed at a nonzero level in at least 75% of samples were considered in this analysis. We retrieved additional transcriptomics data (TCGA RNA-Seq, GSE29623, GSE44076, GSE17536, GSE113513) alongside the previously mentioned multiomics data for our downstream prognostic and diagnostic studies. The TCGA datasets were sourced from the Genomic Data Commons Data Portal [18], while the microarray datasets were retrieved from the NCBI Gene Expression Omnibus (NCBI GEO [19]) database. Differentially expressed genes (DEGs) with a statistically significant pvalue < 0.01 and log2FoldChange (log2FC) were identified using 50 paired normal and CRC samples from the TCGA datasets using DESeq2 methods [20]. The log2FC values were used to quantify the changes in gene expression levels from normal to CRC state. The Robust Multi-array Average (RMA) [21] method was used to normalise the microarray datasets. Detailed information regarding the transcriptomics data utilised in this study can be found in Table 2.

**Table 1.** Information of the CRC multi-omics data obtained from the cBioPortal database.

|  | CNA | mRNA | Mutation |
| --- | --- | --- | --- |
| No. of samples | 616 | 382 | 223 |
| No. of genes | 24776 | 20531 | 12823 |

**Table 2.** A detailed information of the transcriptomics data used in this study. We note the purpose for which each dataset was used in our work. For those datasets which contains no normal samples we denote it with a ‘-’ symbol referring to either ‘none’ or ‘unknown’.

| Dataset | Objective | Number of normal samples | Number of CRC samples | References |
| --- | --- | --- | --- | --- |
| cBioPortal (Firehose Legacy) | Potential prognostic and diagnostic key gene identification | - | 382 | [16] |
| GSE44076 | Diagnostic performance | 98 | 98 | [22] |
| GSE29623 | Prognostic performance | - | 65 | [23] |
| GSE17536 | Prognostic performance | - | 177 | [24] |
| GSE113513 | Diagnostic performance | - | 177 | [25] |
| TCGA-COADREAD | DEGs and Diagnostic performance | 51 | 644 | [18] |

### 2.2 Identification of candidate genes (CGs)

The CNA and mRNA data from the same patient ID were considered for this study to understand the relationship between genetic alteration and gene expression [26]. Our aim here is to identify genes where one type of alteration (either amplification or deletion) is significantly more frequent. First, for each gene, a binary profile is created based on the CNA data in which each gene is assigned a value of 1 if it exhibits genetic alterations (either amplification or deletion) and 0 if it is of the wild type (no copy number change). For each gene, the dominant alteration type (amplification or deletion) is identified using a Binomial test. Let *N* be the total number of subjects (or samples) with CNA, which is the sum of those with amplifications (*A*) and deletions (*D*) that is, *N* = *A* + *D*. To determine the dominant alteration type, the binomial distribution with *N* trials and a success probability of *p* was used. We consider the null hypothesis (*H*_0_) as: The probability of an alteration type being amplification or deletion is equal (*p* = 0.5); while the alternate hypothesis (*H*_1_) as: One alteration type is over-represented (*p* > 0.5) [9]. Now, let *L* = |*A* − *D*|. The objective here is to establish the rejection region where *L* ≥ *ℓ*, where *ℓ* > 0 being a threshold. Since *L* = *A* − (*N* − *A*) = 2*A* − *N*, the probability expression for *L* >= *ℓ* can be represented as:

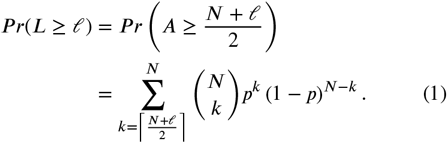

The genes exhibiting the dominant alteration types with significant *p*-value < 0.05 were selected for further analysis.

The mutation information was integrated with the CNA, and the genes altered in at least 10% of the samples were considered for downstream analysis. A one-sided Wilcoxon rank-sum test was performed between the altered and wildtype groups for each gene to determine the effect of the copy number alteration at their mRNA expression level. The genes with concordant changes between CNA and mRNA expression, such as amplification with up-regulated gene expression and deletion with down-regulation in CRC state, were screened. These screened genes were then compared with those that showed differential expression from normal to CRC states. In this study, the genes showing consistent expression changes from normal to CRC state and in between altered and wild-type groups were considered as the “candidate genes” or CGs. These candidate genes were further analysed to understand their significance in the CRC progression.

### 2.3. Construction of WGCN and finding hub genes

In biological systems, genes do not work alone, but they interact with each other to perform a biological function. These interactions between genes also play a significant role in developing complex diseases such as cancer [27]. Weighted gene co-expression network (or WGCN) is a technique in systems biology to explore relationships among genes based on their expression patterns in several samples [28]. WGCN represents a network of genes where each node in the network is a gene and an edge between two nodes in the network indicates the strength or magnitude of their co-expression. WGCN analysis (or WGCNA) emphasises these co-expression relationships among genes and helps identify modules or clusters of highly interconnected genes. These clusters often consist of functionally related genes that can show significant association with the underlying clinical traits. We constructed WGCNs using the candidate genes (CGs) obtained in the previous step to identify the key modules associated significantly with CRC clinical traits (early and late stage). For this analysis, the gene expression data of the CGs was sourced for paired normal and tumour samples from The Cancer Genome Atlas (TCGA) database. Firstly, the goodSamplesgenes function within the WGCNA library [28] was used to check the quality of the genes and samples, followed by hierarchical clustering. Correlations between genes were calculated using Pearson’s correlation coefficient and transformed into a weighted adjacency matrix using the formula, *a*_*ij*_ = |*c*_*ij*_| ^*β*^, where *a*_*ij*_ is an adjacency between gene *i* and gene *j, c*_*ij*_ is Pearson’s correlation, and *β* is the soft-power threshold. Then, the adjacency matrix was transformed into the topological overlap matrix (*T OM*) and average linkage hierarchical clustering was used on the dissimilarity *T OM* (that is, 1 − *T OM*) to identify the modules. The modules with high similarity were merged together (mergeCutHeight = 0.25), and module eigengene (ME) were identified using hierarchical clustering. ME is the first principal component of a given module and represents the the gene expression profiles in a module. The correlation coefficient was calculated between the identified ME and the clinical traits (Normal, Early-stage and Late stage CRC) and a p-value of less than 0.05 was used to screen the key modules. Gene significance (GS) and module membership (MM) were calculated for each gene to identify the hub genes. GS represents the correlation between the individual genes and clinical traits, whereas MM correlates the gene expression profile with the ME. The genes with GS > 0.4 (threshold)and MM > 0.8 (threshold) are considered “hub genes” in our analysis.

### 2.4 Functional enrichment analysis

To understand the biological significance of the identified CGs and modules, we performed the gene ontology (GO) and pathway enrichment analysis using the DAVID database (The Database for Annotation, Visualisation and Integrated Discovery [29]) and Metascape [30]. The GO includes three categories: biological process (BP), molecular function (MF) and cellular component (CC). For pathway enrichment, the KEGG (Kyoto Encyclopedia of Genes and Genomes [31]) database was chosen for the pathway analysis. We also assessed the association of cancer hallmarks with the identified pathways and CGs using the information from the Cancer Hallmark Genes (CHG) database [32].

The regulation of gene expression is essential for maintaining specific cell states; disruption in this regulation can lead to various diseases. Gene expression is controlled by various factors, including transcription factors (TFs) and miRNAs, which regulate expression at the transcriptional and post-transcriptional levels, respectively [33]. Therefore, to understand the regulatory mechanism involved in the regulation of the identified CGs, TFs and miRNA were predicted for the CGs using g:GOSt within the g:Profiler tool [34, 35].

### 2.5. Screening of key genes for prognosis

To analyse the impact of the identified CGs and the hub genes in the progression of CRC, we integrated the clinical information, such as overall survival (OS), with the mRNA expression for each sample. Samples missing the OS status of the patients were excluded from the study, and a total of 368 samples were analysed for survival analysis. Firstly, univariate Cox proportional hazards regression analysis was used to screen for genes significantly associated with CRC prognosis. For this, we used two R packages, survival [36] and survminer [37] with a significance threshold of p-value less than 0.05 (*p* < 0.05). After that, the Least absolute shrinkage and selection operator (LASSO) Cox method with 10-fold cross-validations was used to control the collinearity between genes using the glmnet R package [38]. The λ value in which mean square error is minimum was selected as the optimum λ parameter. Genes whose coefficients were not zero were considered strongly related to survival. Then, Multivariate Cox proportional hazards regression analysis was conducted to identify the key genes involved in the progression of CRC [39]. These genes were further used to construct risk scores to develop a prognostic model to predict the prognostic status. The risk score was calculated for each sample by using the following formula:

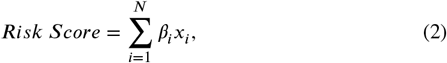

where *N* is the total number of genes, *x*_*i*_ is the expression levels of gene *i* in a particular patient sample and and *β*_*i*_ denotes the coefficient (meaning, weight) for gene *i* in the multivariate Cox regression analysis. The risk score was calculated for each patient sample and then categorised into high-risk and low-risk groups based on the median value. Then, Kaplan–Meier survival curves [40] were plotted to assess the ability of these genes to distinguish patients in the low-risk group from those in the high-risk group, with a statistically significant *p*-value. The risk score calculation was also done on two independent validation datasets, GSE17536 and GSE29623, where each patient sample was assigned a risk score. Furthermore, the predictive performance of this risk score model was evaluated over different time intervals, specifically at 1, 3, and 5 years, using a time-dependent receiver operating characteristic (ROC) curve analysis. The effectiveness of the risk score model in predicting patient outcomes was evaluated using the ROC curve analysis on independent validation datasets GSE17536 and GSE29623.

### 3.6. Identification of diagnostic biomarkers

The genes that were identified as “hub” genes in late-stage CRC through WGCNA were selected for the diagnostic classification of normal and CRC samples. Among these hub genes, six were common hub genes across both the early and late stages of CRC. The TCGA RNA-Seq data, GSE44076 and GSE113513 datasets with normal and CRC samples were used for this analysis. In this study, we used some classical statistical and machine learning (ML) methods available within the Scikit-learn library [41] for our analysis. Specifically, we used two unsupervised methods, principal component analysis (PCA) and t-distributed stochastic neighbour embedding (t-SNE), to visualise the data. We used two kinds of supervised machine learning (ML) techniques (a) Logistic Regression, a linear model, and (b) Random Forest, an ensemble of trees to evaluate the diagnostic efficacy of the seven identified hub genes in distinguishing between normal samples and CRC samples. Here, the seven genes with their expression values serve as the “input features” for the two machine learning models.

Evaluating two different kinds of ML models could provide a statistically more robust and reliable estimate of the genes’ diagnostic efficacy. To ensure a strong regularisation, we used *L*_2_ penalty and inverse regularisation strength of *C* = 0.1 for training our Logistic Regression model. This allows the model to focus on specific genes that are much more diagnostic towards the CRC status. We used 50 trees in our Random Forest model. For both these models, all other hyperparameters were left to their default values within the Scikit-Learn library. Furthermore, to ensure the comparability between the training and validation data, min-max normalisation, available within Scikit-Learn, was applied to the external validation datasets using the same scaling parameters derived from the training dataset. We note, however, that while such a simple normalisation works in this study, it may not necessarily be applicable to other datasets. This is a long-standing issue with the analysis of RNA-seq datasets using ML [42], which poses a classic challenge of distribution shift [43]. We use several evaluation metrics, such as accuracy, sensitivity, specificity, precision, recall, ROC-AUC and Matthews correlation coefficient (MCC), to evaluate our trained models on testing datasets. For this experiment, we run the ML modelling with several random seeds to get an average estimate of the predictive performance. The relevant details are provided in the Results.

## 3. Results

### 3.1. Candidates genes in CRC

We integrated the CNA information with the mRNA data for 376 samples, and the results showed the effect of the copy number on the gene expression, as shown in Figure 2A. We observed an increase in the *z*-score of the genes with gain or amplification compared to the gene with the loss and deletion CNA. A total of 7571 genes were screened, showing dominant alteration with concordant changes in their gene expression between the altered and wild-type groups. For example, we observed that the dominant alteration type for the MYC gene was amplification and up-regulated mRNA expression was observed in the altered group compared to the wild-type group. Similarly, for APC genes, the dominant alteration was deletion and expression in the altered group observed was down-regulated (Figure 2B). The APC gene is frequently mutated in colorectal cancer (CRC) and is recognised as a crucial tumour suppressor gene. Conversely, MYC is identified as an oncogene, contributing to the oncogenic process [44]. These findings strongly align with our research, showing the significance of integrating genetic alteration information with mRNA. *LRRC69* gene (Figure 2B) with amplification CNA and up-regulated gene expression in CRC state was identified as a key gene impacting CRC patients’ survival. These genes were further screened by mapping their CNA with the identified DEGs, such as amplification with up-regulated DEGs and deletion with down-regulated DEGs. Based on this criteria, 3028 genes were screened as candidate genes (CGs) containing 1968 up-regulated genes with amplification CNA type and 1060 down-regulated genes with deletion CNA type. Our study found the *MYC* and *APC* to be up-regulated and downregulated, respectively, in the CRC.

**Figure 2:**
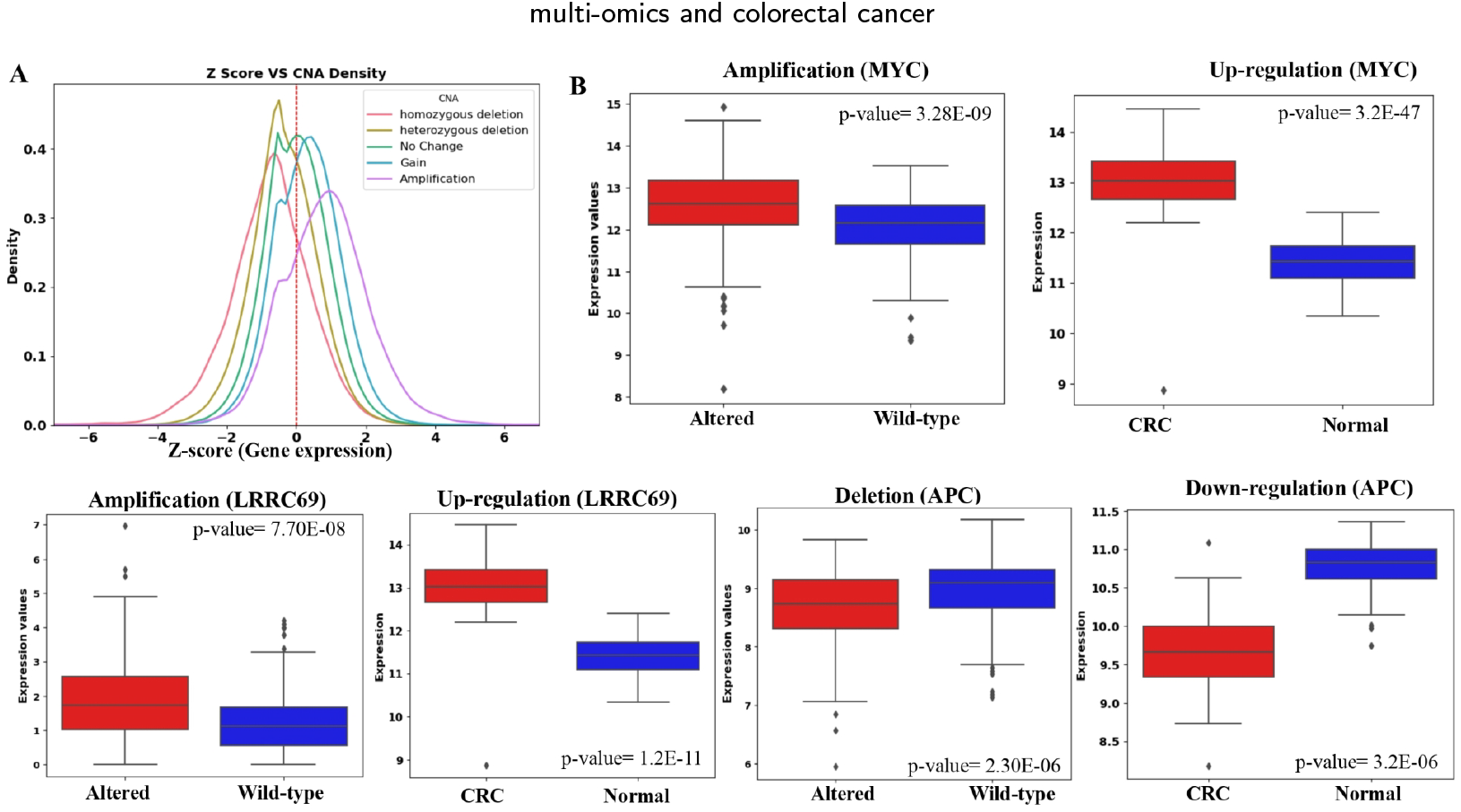
Integration of CNA with gene expression data. **(A)** Density plot showing the association of *z*-score (gene expression, Yaxis) versus copy number alteration (X-axis) across 376 samples of CRC. **(B)** Box plots represent the expression of *MYC, LRRC69* and *APC* genes in the altered and wild-type (No genetic alteration) groups and the Normal and CRC groups, respectively. *p*-value in each of the box plots represents the statistical significance between the gene expressions for the samples in the two considered groups.

### 3.2. Identification of clinically significant modules and hub genes

Based on the quality check of the genes and samples, two samples were removed from the analysis (Figure 3A-B). We used the soft-threshold power, *β* = 18 (scale-free *R*^2^ = 0.8) to construct a scale-free network, and co-expression modules were identified by hierarchical clustering (Figure 3C-E). Seven modules, including blue (306), brown (245), green (51), grey (1888), red (34), turquoise (337) and yellow (61), were identified in this analysis. The number in brackets are representing the number of genes in the respective module. After identifying the modules, the correlation between each module and clinical traits was analysed. It was found that the green and red modules showed a significantly high positive correlation, while the blue and turquoise modules showed a significantly high negative correlation with early and late-stage traits. As a result, these four modules were selected as key modules for further study (Figure 3F).

**Figure 3:**
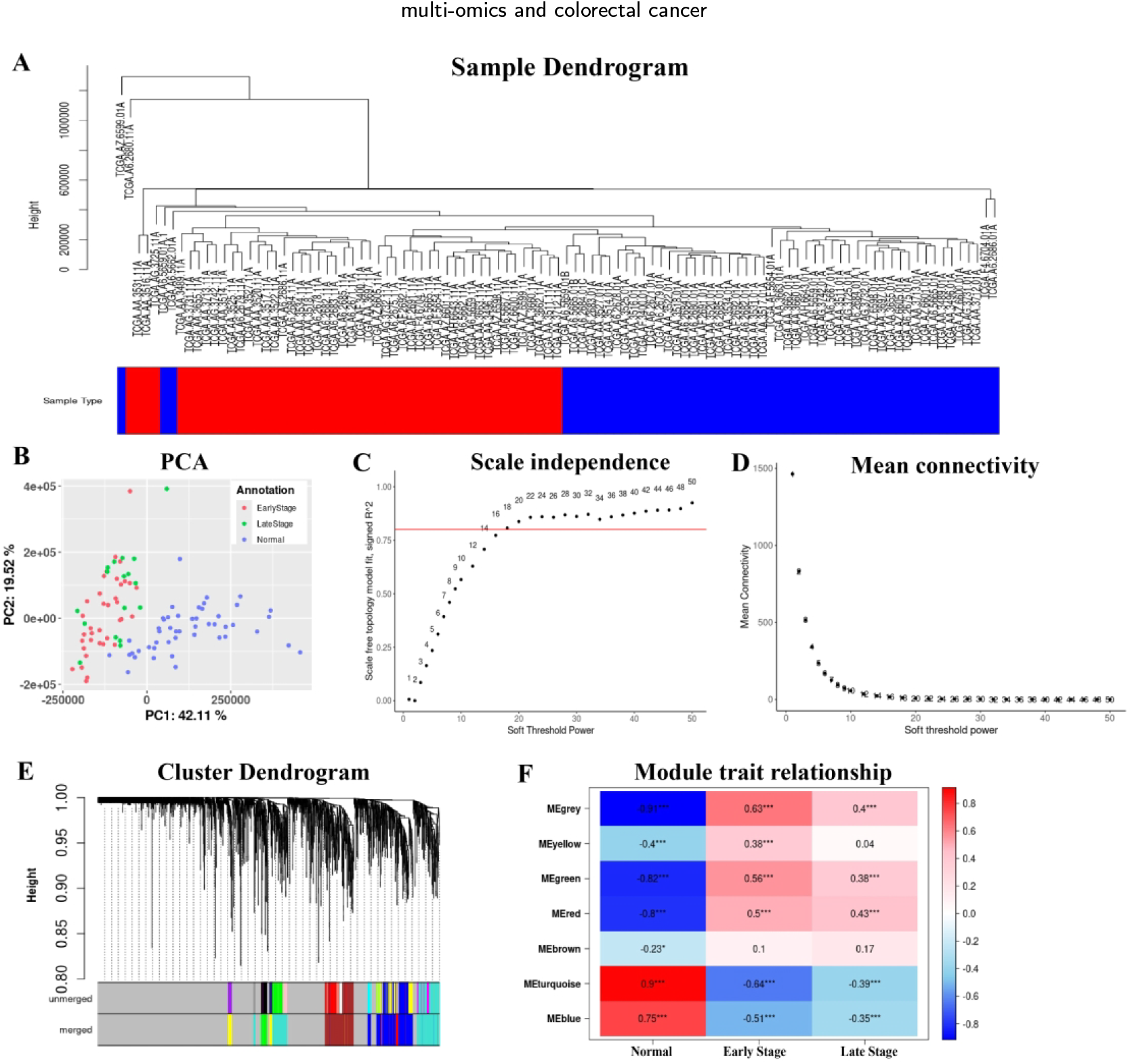
WGCNA analysis of the candidate genes. **(A)** Samples dendrogram and traits heatmap to detect the sample outliers. The clustering was based on the expression data of candidate genes between normal samples and CRC samples. The blue and red colours represent the CRC and normal samples, respectively. **(B)** PCA analysis of the selected samples. **(C)** Screening of soft-thresholding powers. **(D)** Analysis of mean connectivity for various soft-thresholding powers. **(E)** Dendrogram clustered of the CGs based on a dissimilarity measure (complemented topological overlap matrix, 1 − *T OM*) before and after merging. **(F)** Heatmap representing the correlation between the module eigengene and clinical traits. Here ‘*’: *p* < 0.01 and ‘***’: *p* < 0.001.

Gene significance and module membership were plotted for the selected modules to identify the hub genes associated with both the early and late stages of CRC, as shown in Figure 4. Seven hub genes, including *DSCC1* from the green module, *ETV4, KIAA1549* from turquoise, *NOP56, RRS1*, and *TEAD4* from red, and *ANKRD13B* from the blue module, with GS > 0.4 and MM ≥ 0.8 were found to be significant in the association of CRC.

**Figure 4:**
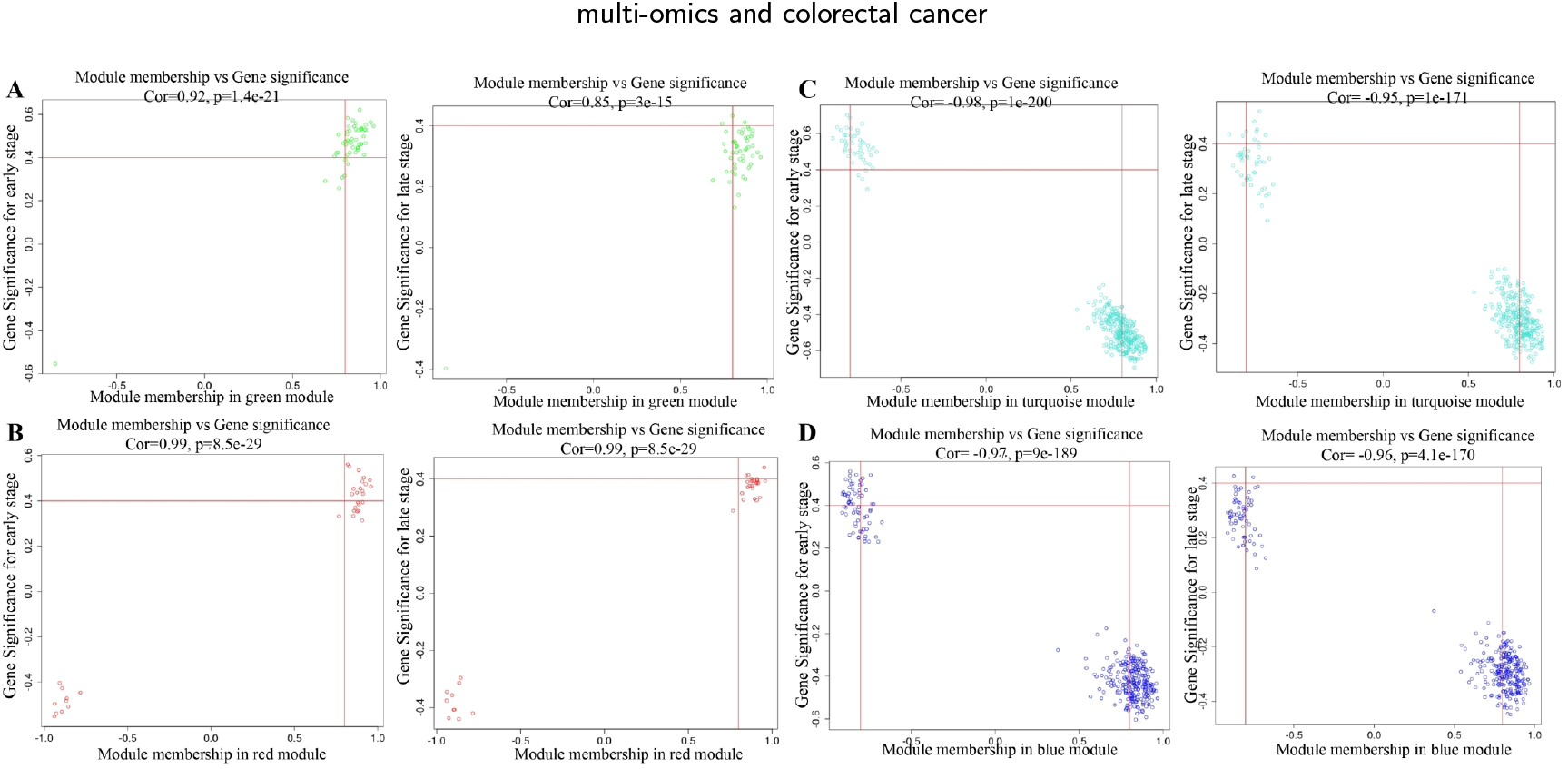
Scatter plot representing the correlation between the selected modules and gene significance for the early and late stage in **(A)** Green **(B)** Red **(C)** Turquoise and **(D)** Blue, respectively. Cor and p denote the correlation coefficient and p-value, respectively.

### 3.3 Functional annotation of identified CGs and modules

The biological significance of the CGs and modules was explored through GO and KEGG Pathway analysis. The analysis revealed that the CGs were mainly enriched in various biological processes, including transcriptional regulation, cell cycle, apoptosis, protein ubiquitination, and protein phosphorylation (Figure 5). As for cellular components, CGs were mainly found to be predominantly located in the nucleus, cytosol, cytoplasm, and nucleoplasm, whereas molecular function analysis showed enrichment in protein binding, metal ion binding, RNA binding, and ATP binding (Figure 5). The KEGG Pathway enrichment analysis showed that CGs were mainly involved in the endocytosis, cell cycle, spliceosome, wnt signalling and mTOR signalling pathways as presented in Table 3. The Metascape results revealed significant associations between the enriched pathways, primarily focusing on metabolism and cell cycle-related processes (Figure 6. The identified CGS genes were crossreferenced with established CRC-associated genes and oncogenic driver genes using the DisGeNET [45] and OncoVar [46]databases, respectively. This comparison revealed that 29 genes overlapped with the known CRC-associated gene, 21 driver genes, and 116 genes with oncogenic mutation datasets (Table S1). These findings highlight the significance of integrating multi-omics approaches, as it enhances the identification of essential genes involved in the progression of cancer. The tissue specificity enrichment analysis revealed that CGs are primarily enriched in the gastrointestinal system tissues, including the colon, duodenum, rectum, small intestine, and stomach (Figure 7A).

**Table 3.**
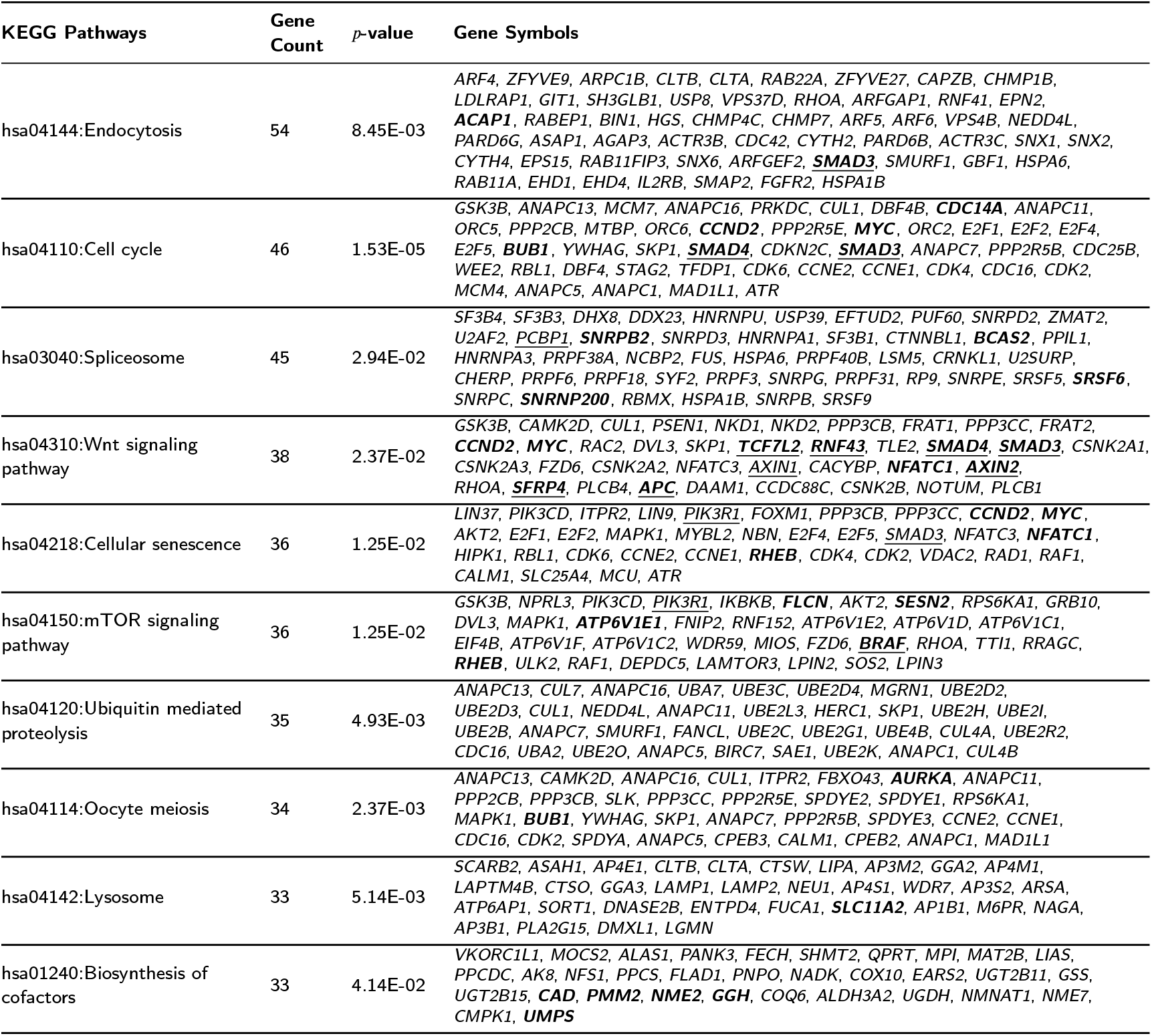
Top 10 KEGG Pathways for the identified CGs based on the maximum number of genes and *p*-value. The gene symbols highlighted in **bold** represent known CRC-associated genes based on the curated DisGeNET database [45] and the genes with **underline** are the driver oncogenes for CRC as per the OncoVar database [46].

**Figure 5:**
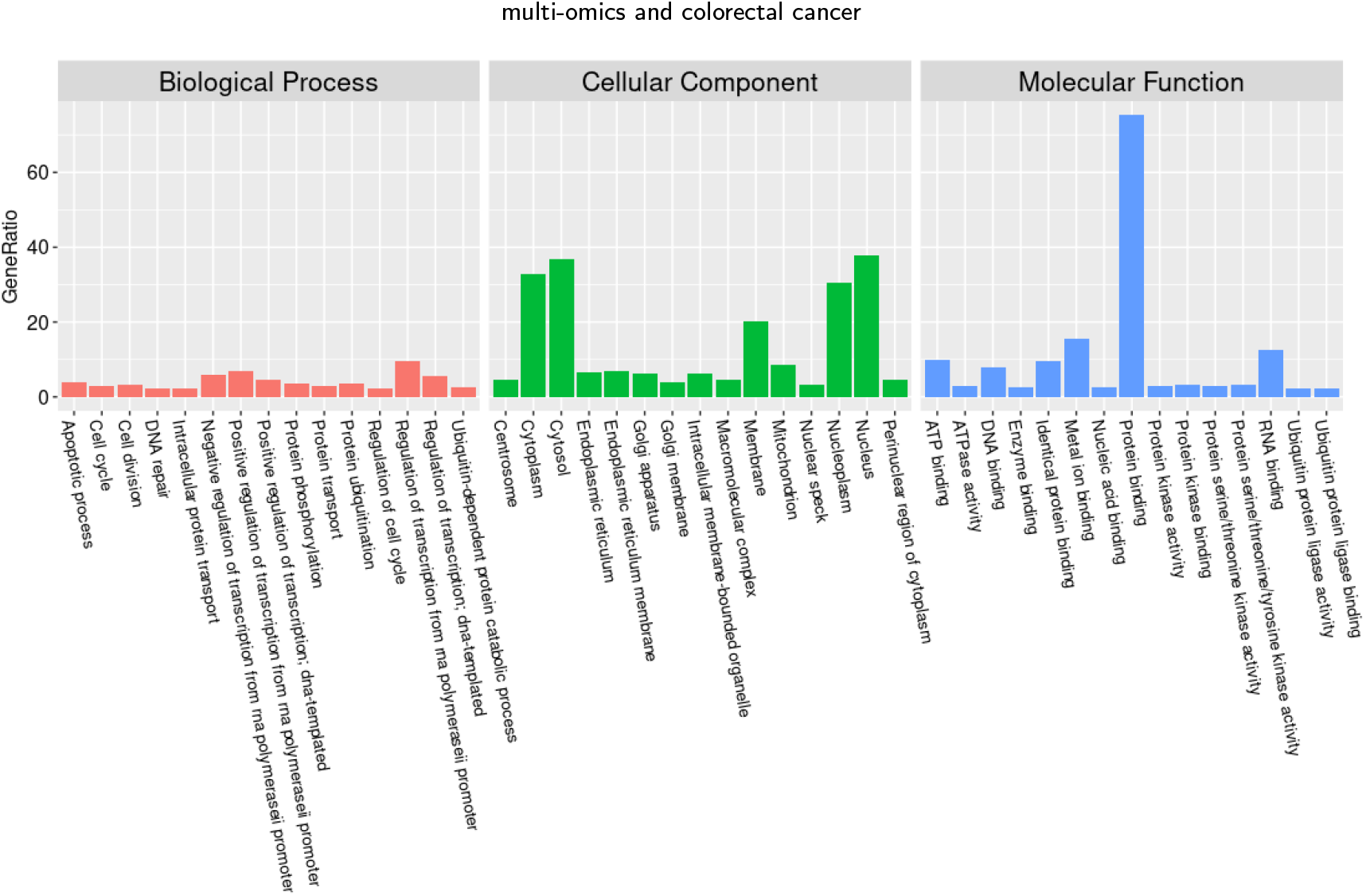
Gene ontology analysis of the identified CGs. Bar graph representing the top 15 biological processes, cellular components and molecular functions.

**Figure 6:**
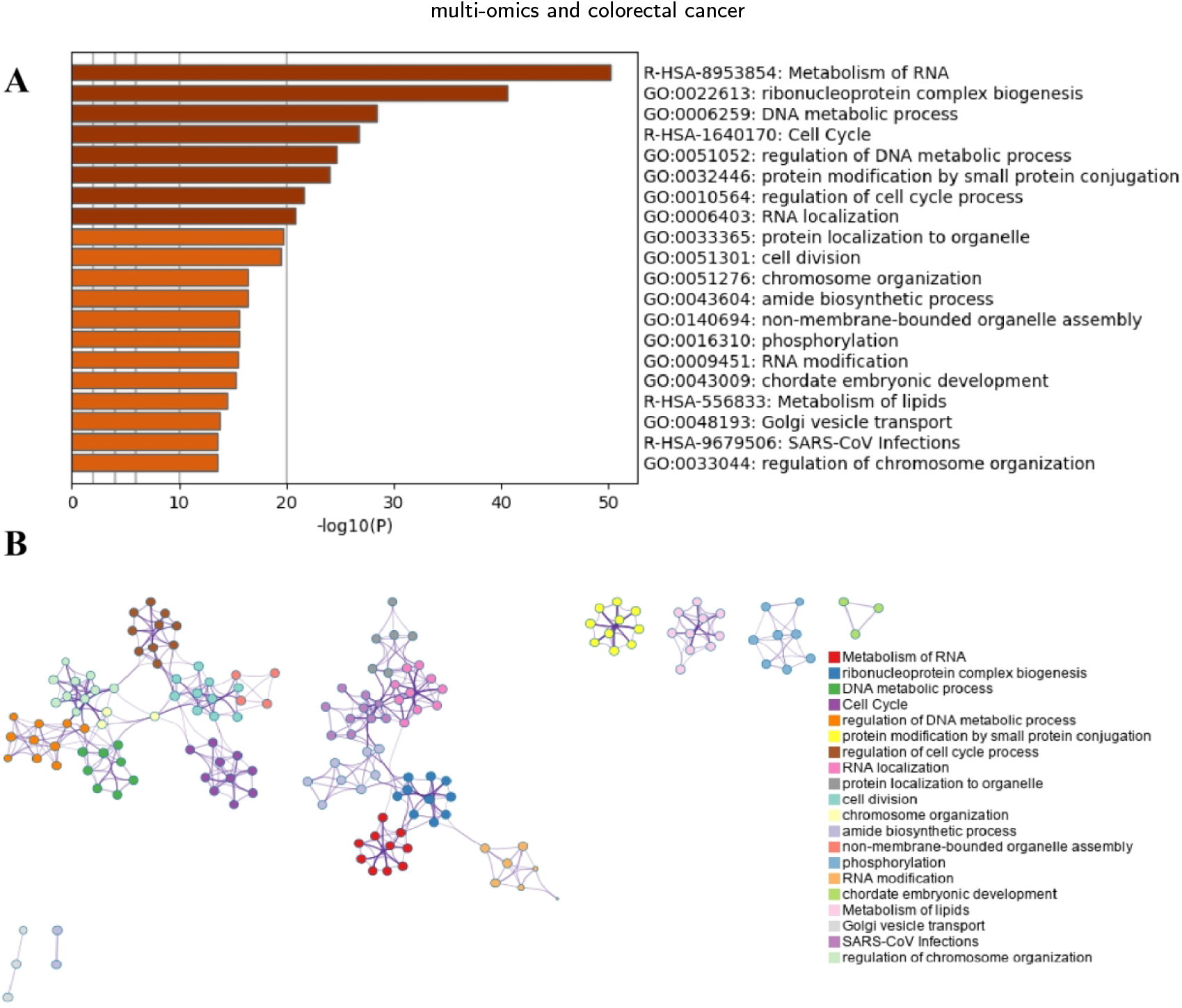
Enrichment analysis. **(A)** Bar graph representing the top enriched clusters, and **(B)** Network representing the associations between the enriched clusters.

**Figure 7:**
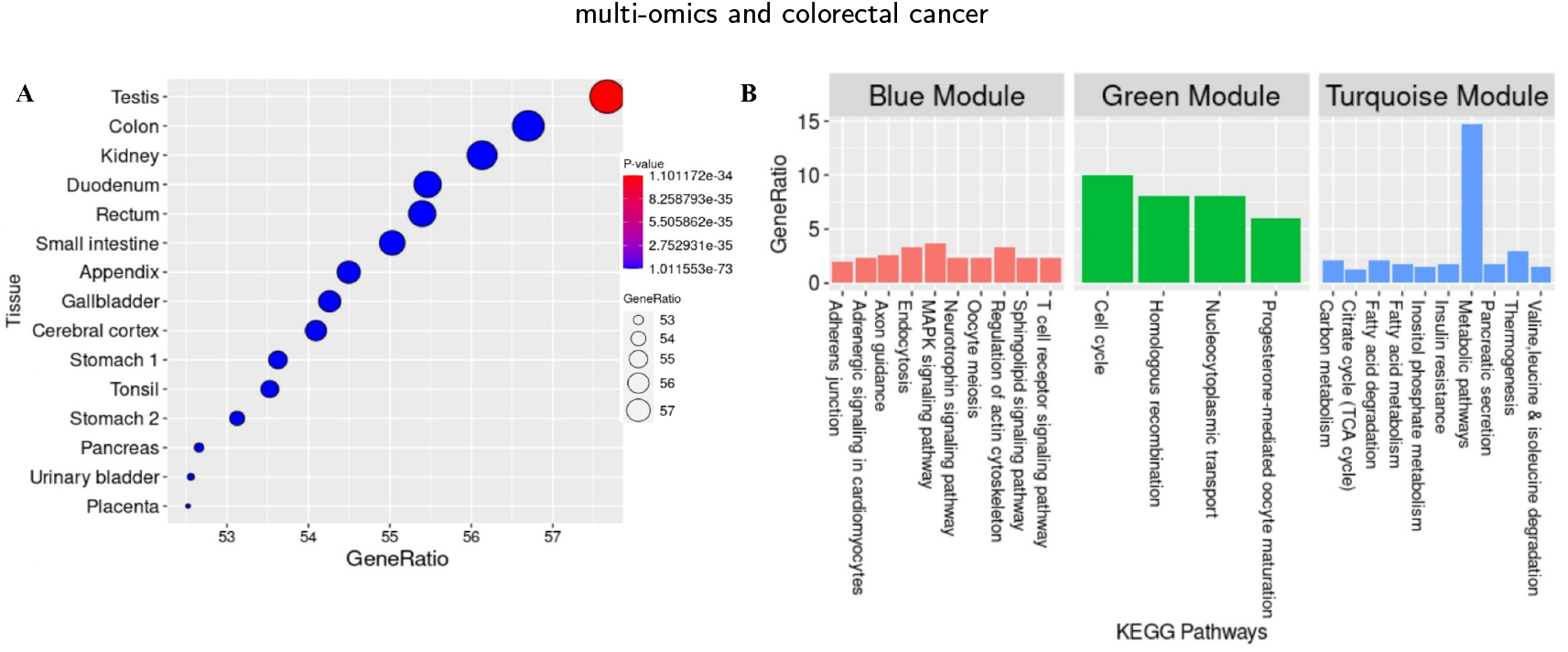
Functional enrichment analysis. **(A)** Dot plot representing the tissue-specific enrichment analysis of CGs. **(B)** KEGG Pathway enrichment analysis of the identified key modules.

The pathway enrichment analysis was also performed for the identified modules, and the results revealed that the blue module was mainly enriched with signalling pathways such as T cell receptor signalling, sphingolipid signalling, and MAPK signalling, whereas the green module consisted of cell cycle-related pathways and turquoise modules involved in the metabolic-related pathways (Figure 7). We did not find any significant pathways for the red modules. For the identified pathways and CGs, we also assessed their association with the cancer hallmark and 13 pathways and 428 CGs were found to be associated with at least one type of cancer hallmark (Figure 8). The TGF-*β* signalling pathway was found to be associated with the five cancer hallmarks, including Activating Invasion and Metastasis, Enabling Replicative Immortality, Evading Growth Suppressors, Resisting Cell Death and Sustaining Proliferative Signaling. The Wnt signalling pathway was also associated with three hallmarks: Activating Invasion and Metastasis, Enabling Replicative Immortality and Sustaining Proliferative Signaling cancer hallmarks. Among the identified hub genes, the *TEAD4* gene was found to be associated with the Sustaining Proliferative Signaling and Evading Growth Suppressors cancer hallmarks. These findings suggest that the identified CGs, modules and hub genes may play a crucial role in the progression of CRC and deepen our understanding of their significance.

**Figure 8:**
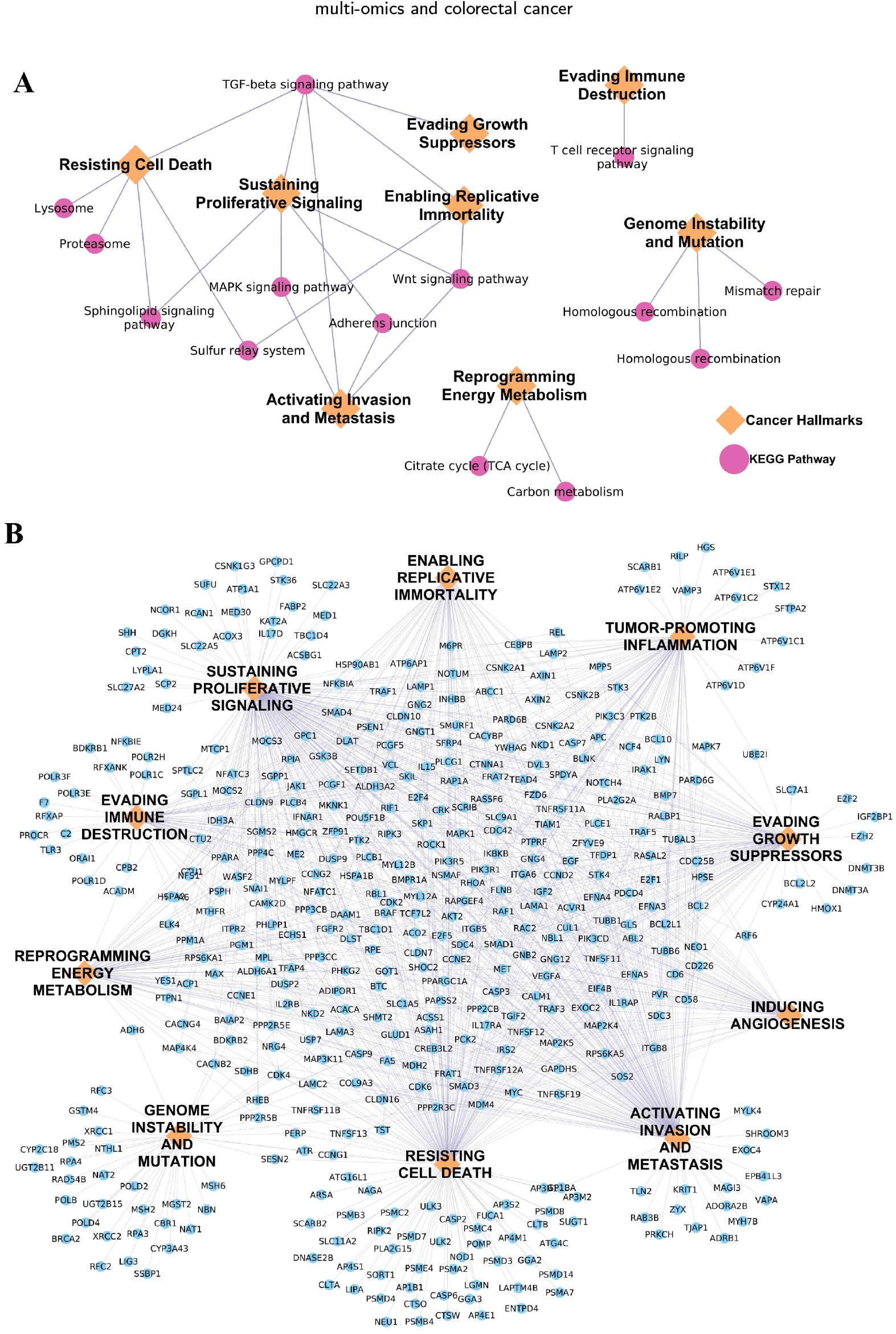
Association of cancer hallmarks with the **(A)** identified KEGG Pathways and **(B)** candidate genes.

### 3.4 Analysis of gene regulatory networks

Gene regulatory networks consisting of the TF-mRNA and miRNA-mRNA were constructed to understand the regulatory mechanisms of the identified CGs. A total of 20 miRNAs were found to regulate the 1607 genes, whereas 229 TFs were regulated at the transcription level (Table S1). Among 229 TFs, 26 were observed to be CGs of our analysis (Figure 9A), which includes *ETV4*, one of the hub genes identified in this study.

**Figure 9:**
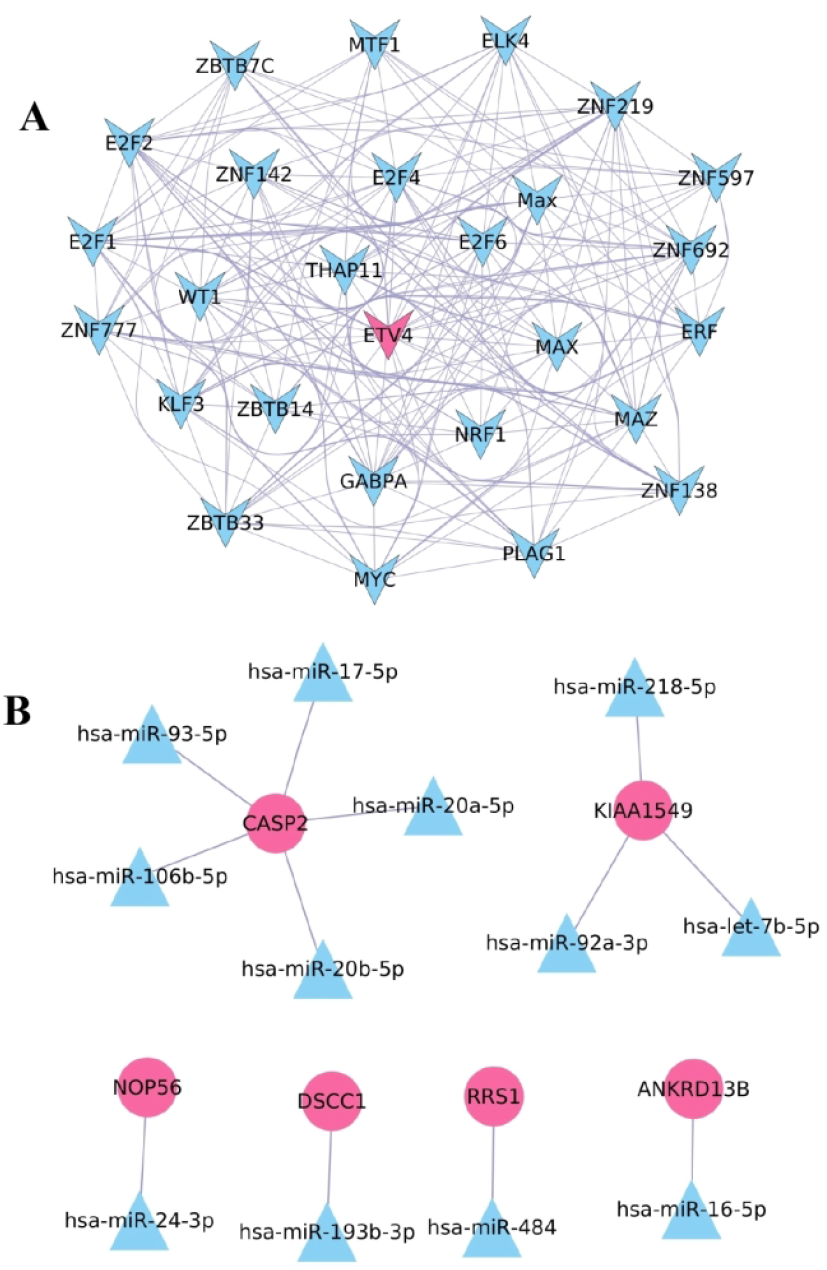
Gene regulatory network analysis. **(A)** Transcription factors identified as CGs in our analysis, with *ETV4* identified as the hub gene. **(B)** miRNA-gene network, representing the regulation of key and hub genes by miRNA. The magenta colour represents the key and hub gene.

### 3.5. Identification of key genes in the progression of CRC

The mutations or genetic alterations of genes associated with cancer progression are known as driver genes in cancer [47, 48]. In this study, we analysed the impact of identified CGs on the survival of CRC patients and defined them as key driver genes in CRC progression. For this, univariate Cox proportional hazards regression analysis with Lasso regression and multivariate Cox proportional hazards regression analysis were done to screen key driver genes. The univariate Cox proportional hazard regression analysis identified 383 genes with a significant (p-value < 0.05) effect on the progression. These 383 genes were then subjected to LASSO regression, and *λ* was determined, where partial likelihood deviance was the smallest. For *λ* = 0.0311, genes with coefficients greater than 0 were screened and used for multivariate Cox proportional hazards regression analysis (Figure 10A-C). A total of four genes, *CASP2, HCN4, LRRC69* and *SRD5A1*, were obtained from multivariate analysis with a hazard ratio greater than 1 to calculate the risk score for each patient. The risk score was calculated using the linear equation: (0.844· *CASP2* +0.433· *HCN4* + 0.264 · *LRRC69* + 0.505 · *SRD5A1*). We observed that the prognosis of the low-risk group was significantly better than that of the high-risk group of CRC patients (Figure 10D). The risk scores calculated for the external validation datasets GSE29623 and GSE17536 consistently showed higher survival probabilities in the low-risk group compared to the high-risk group, showing the significance of the risk score in predicting patient outcomes (Figure S1). The risk score-based prediction model analysis using time-dependent ROC showed predictive accuracy, with AUC values of 0.90 for one year, 0.92 for three years, and 0.85 for five years. The risk score-based prediction model was validated using external datasets, revealing AUC values of 0.70, 0.71, and 0.69 for GSE29623 and 0.68, 0.63, and 0.63 for GSE167536 at 1, 3, and 5 years, respectively. This validation confirms the model’s predictive accuracy over different time periods in distinct datasets (Figure 10E-G). These results suggest that the risk-based prediction model used in this study has potential clinical value as a prognostic tool for predicting patient survival.

**Figure 10:**
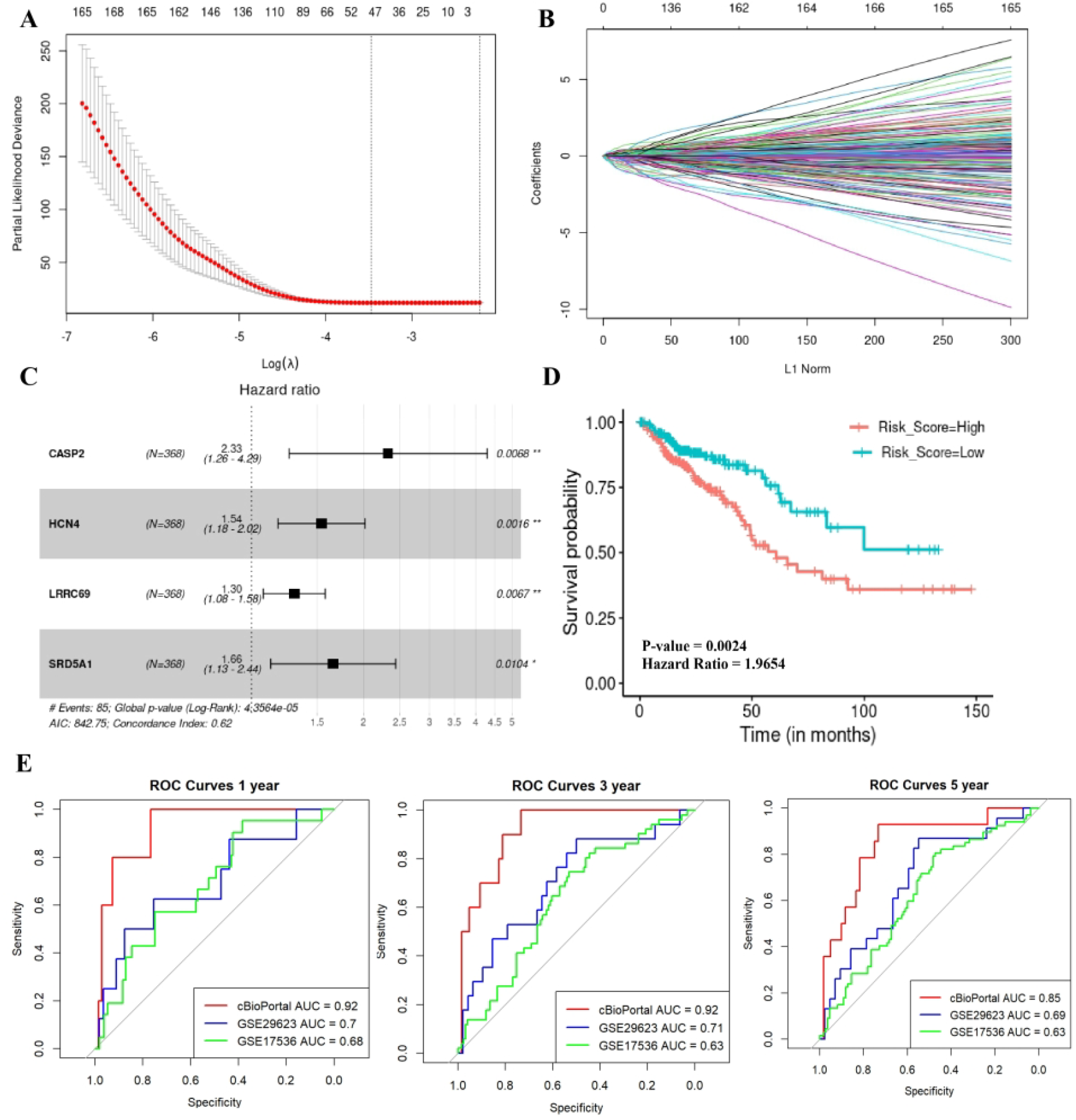
Prognostic significance of identified CGs. **(A)** The LASSO Cox regression model to identify optimal λ value. **(B)** LASSO coefficient profiles of prognostic genes. **(C)** Forest plot for multivariate Cox regression of selected four genes. **(D)** Survival curve for CRC patients with low-risk (Risk Score < median value) and high-risk (Risk Score > median value) groups. **(E)** ROC curves for the model representing 1-year, 3-year and 5-year patient survival prediction for the cBioPortal datasets, GSE29623 and GSE17536. * representing the p-value.

### 2.6. Identification of diagnostic biomarkers for CRC

The results from PCA and t-SNE analyses revealed that the seven selected hub genes could differentiate between normal samples, early stage and late-stage CRC samples as shown in Figure 11. Therefore, we used the expression of these seven genes to create a machine learning (ML) model that can accurately predict the CRC status. As described in Methods, we used two classic machine learning (ML) techniques, such as Logistic regression and Random Forest to construct these CRC diagnostic models using these seven hub genes and their expressions as input features. In what follows, we study the diagnostic performance of these ML models in two scenarios: (1) Predicting normal vs CRC samples and (2) Predicting early-stage vs late-stage CRC.

**Figure 11:**
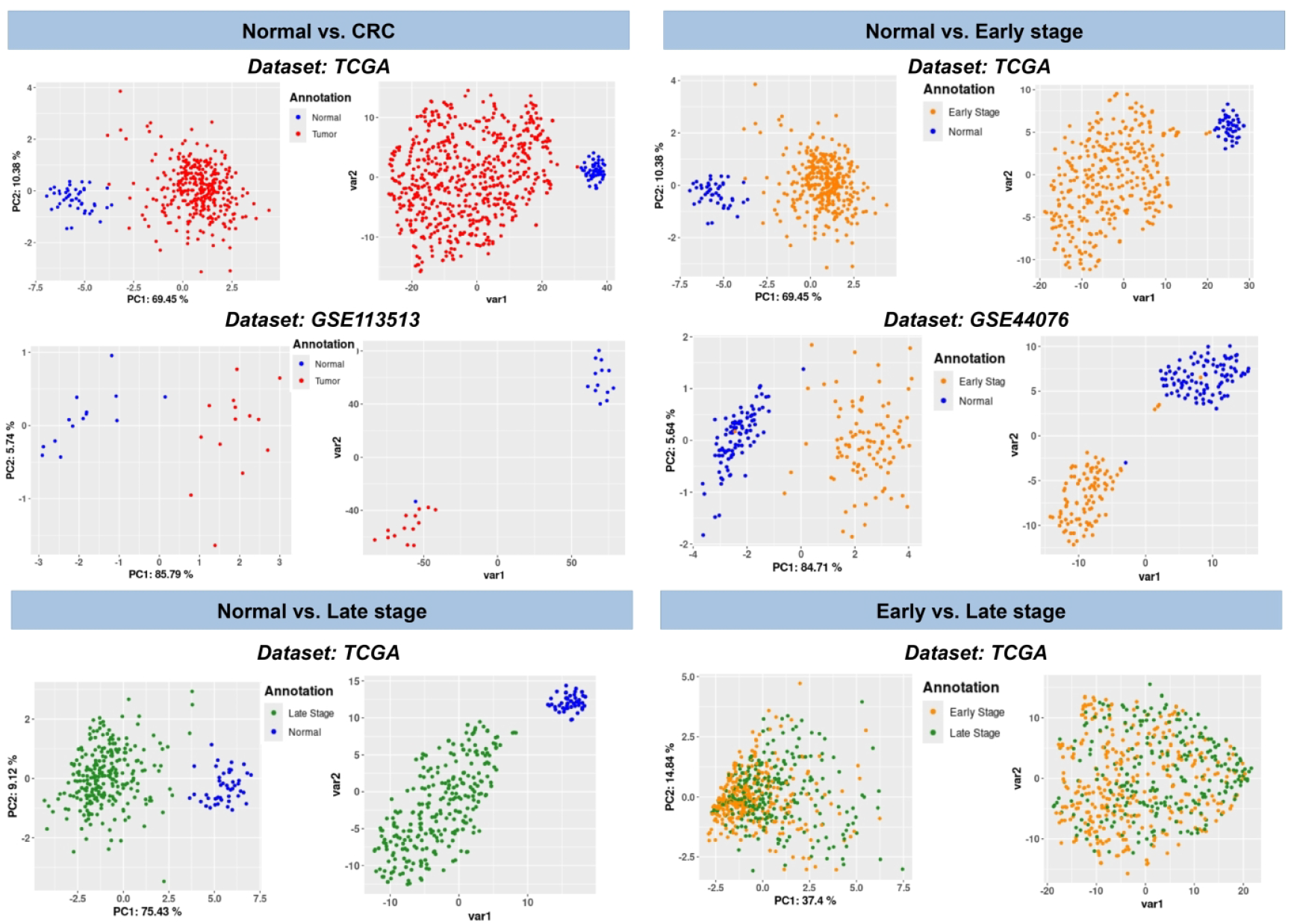
A visual analysis of the diagnostic performance of the seven hub genes using PCA and t-SNE across the TCGA RNA-Seq, GSE44076, and GSE113513 datasets.

#### 3.6.1. Predicting normal vs CRC Samples

The TCGA datasets were used to discriminate the normal samples from all CRC samples, as well as early-stage and late-stage CRC samples. The TCGA dataset was divided into training, test, and internal validation sets using a random split ratio of 60:20:20 with stratified splitting. The ratio of normal and CRC samples was also considered during the splitting process and repeated 100 times to ensure a robust and reliable predictive analysis. Four performance metrics, average accuracy, precision, recall, F1 score and MCC score, were calculated and summarised and are reported here in Table 4. The best-performing model was selected and assessed on both internal and external validation datasets (GSE113513 and GSE44076 for normal vs all CRC and normal vs early-stage CRC), resulting in AUC values of 0.96 and 0.99 with the logistic regression model (Figure 12) and 0.89, and 0.93 with the random forest model (Figure S2, Table S2), respectively. The validation of normal vs. late-stage samples on the internal validation datasets revealed ROC and precision-recall curves with an AUC value of 1. These results highlight the significance of the seven identified hub genes in distinguishing between normal and early-stage as well as normal and late-stage CRC samples.

**Table 4.** Average performance (from 100 different runs) metrics of the Logistic Regression model for predicting CRC status.

| Study | Validation | AUC | Sensitivity | Specificity | Accuracy | Precision | Recall | F1 Score | MCC |
| --- | --- | --- | --- | --- | --- | --- | --- | --- | --- |
| Normal vs. CRC | *TCGA<sub>Test</sub>* | 0.981 | 1.000 | 0.961 | 0.997 | 0.997 | 1.000 | 0.999 | 0.979 |
|  | *TCGA<sub>Validation</sub>* | 0.982 | 1.000 | 0.961 | 0.998 | 0.997 | 1.000 | 0.999 | 0.980 |
|  | GSE113513 | 0.935 | 1.000 | 0.857 | 0.935 | 0.886 | 1.000 | 0.939 | 0.878 |
| Normal vs. Early-Stage | *TCGA<sub>Test</sub>* | 0.994 | 0.999 | 0.989 | 0.998 | 0.999 | 0.999 | 0.999 | 0.990 |
|  | *TCGA<sub>Validation</sub>* | 0.994 | 0.999 | 0.989 | 0.998 | 0.999 | 1.000 | 0.999 | 0.992 |
|  | GSE44076 | 0.990 | 1.000 | 1.000 | 0.990 | 0.990 | 0.990 | 0.990 | 0.980 |
|  | *GSE44076<sub>Test</sub>* | 0.977 | 0.959 | 0.994 | 0.977 | 0.995 | 0.959 | 0.976 | 0.955 |
|  | *GSE44076<sub>Validation</sub>* | 0.982 | 0.938 | 1.000 | 0.982 | 0.989 | 0.976 | 0.982 | 0.965 |
|  | TCGA | 0.989 | 0.938 | 1.000 | 0.981 | 1.000 | 0.978 | 0.989 | 0.920 |
| Normal vs. Late-Stage | *TCGA<sub>Test</sub>* | 1.000 | 0.999 | 1.000 | 0.999 | 1.000 | 0.999 | 1.000 | 0.998 |
|  | *TCGA<sub>Validation</sub>* | 0.999 | 0.999 | 1.000 | 0.999 | 1.000 | 0.999 | 1.000 | 0.998 |

**Figure 12:**
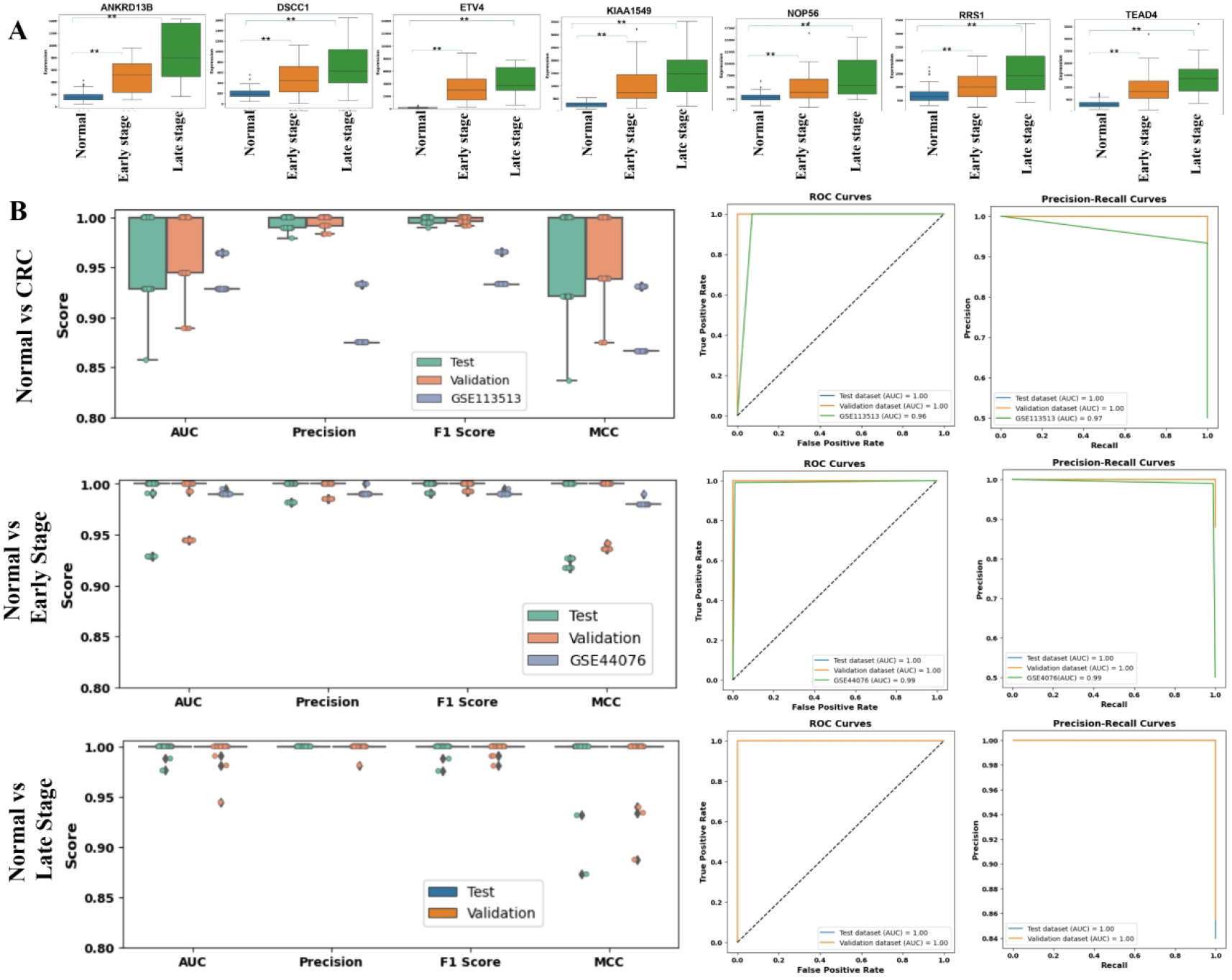
Diagnostic performance of hub genes. **(A)** Boxplots represent the change in expression of seven genes from normal to early stage and late stage CRC. ‘*’ denotes p-value < 0.01; **(B)** represents the diagnostic performance evaluation of predictive models in CRC using internal and external validation datasets, showcasing the respective ROC and precision-recall curves.

#### 3.6.2. Early-stage vs Late-stage CRC

Building a model to predict early-stage vs late-stage CRC samples is a relatively harder problem than predicting normal vs CRC samples, primarily owing to the inherent complexity of CRC. By using only gene expression data, the seven hub genes showed poor performance in discriminating between early and late-stage CRC samples. Therefore, we integrated the gene regulatory information to screen the genes, which might be able to differentiate the early-stage samples from late-stage CRC samples. First, we calculated the regulatory strength of each gene by considering all its interactions with other genes in both early and late-stage CRC samples by using the equation:

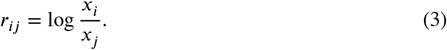

Here, *r*_*ij*_ represents the regulation strength for the gene *i* and *j* connected in the gene regulatory network, *x*_*i*_ and *x*_*j*_ represents the expression value of gene *i* and gene *j* [49]. Then, the cumulative regulatory strength of each gene (*i*) was calculated by taking the sum of all its interacting neighbours by using the equation:

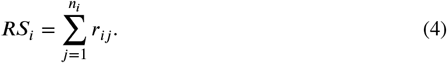

Here, *RS*_*i*_ denotes the cumulative regulatory strength of gene *i, n*_*i*_ is the number of first (or 1-hop) neighbours of gene *i* (interacting genes) in the gene regulatory network. The regulatory strength was calculated for each sample. These regulatory strengths of the genes were used for feature selection using the mRMR algorithm. Among 3028 candidate genes, only 1703 were found to be present in the gene regulatory network. These 1703 genes were then ranked using the mRMR feature selection algorithm, and then the top 5, 10, 15, 20 and 25 genes were selected for further analysis. The analysis using PCA and t-SNE revealed that genes with regulatory strength were unable to differentiate between the early and late stages of colorectal cancer (Figure S3). To improve the predictive accuracy further, the top 5, 10, 15, 20, and 25 genes (Table 5) based on regulatory strength were selected and used to build a predictive model. Comparing the performance of this model with the one based on gene expression, it was found that genes selected for their regulatory strength provided better predictions (Figure 13, Figure S4) and (Table 6). The top 15 genes were chosen for further analysis, and the ROC and precision-recall results showed that these genes performed better in distinguishing between early-stage and late-stage samples.

**Table 5.**
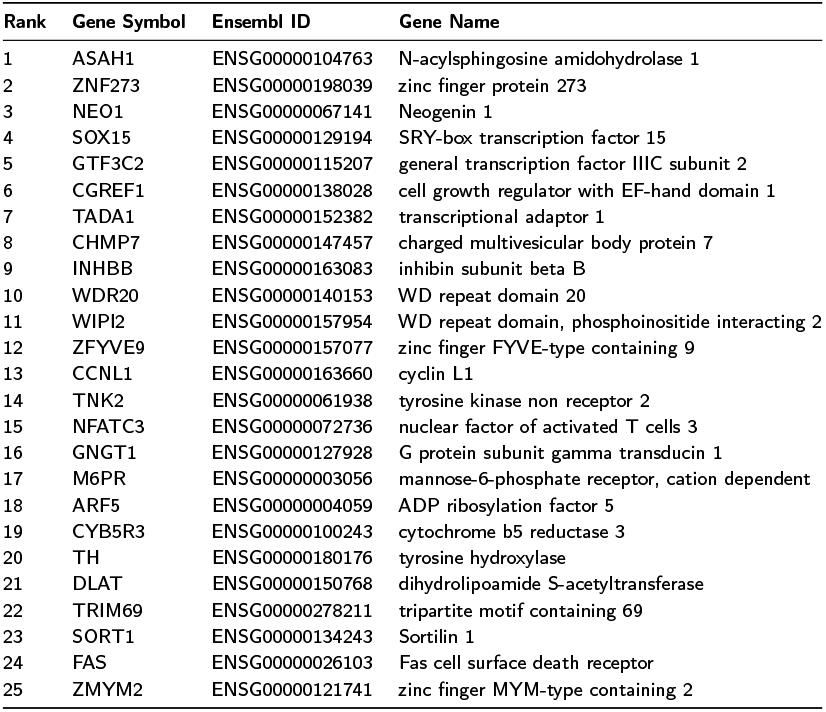
The top 25 genes selected based on their regulatory strength (Eq. (4)). Each gene is provided here with its Ensembl ID.

**Table 6.** Comparison of the predictive performance between gene expression-based (seven hub genes) and regulatory strength-based selected genes (Top 5, 10, 15,20 and 25) using logistic regression to predict early and late-stage status.

| No. of Genes | Validation | AUC | Sensitivity | Specificity | Accuracy | Precision | Recall | F1 Score | MCC |
| --- | --- | --- | --- | --- | --- | --- | --- | --- | --- |
| Seven genes | $TCGA_{Test}$ | 0.551 | 0.297 | 0.804 | 0.580 | 0.553 | 0.297 | 0.381 | 0.120 |
| | $TCGA_{Validation}$ | 0.553 | 0.297 | 0.804 | 0.581 | 0.561 | 0.296 | 0.384 | 0.126 |
| Top 5 | $TCGA_{Test}$ | 0.643 | 0.531 | 0.755 | 0.656 | 0.634 | 0.531 | 0.576 | 0.295 |
| | $TCGA_{Validation}$ | 0.636 | 0.531 | 0.755 | 0.648 | 0.626 | 0.526 | 0.569 | 0.280 |
| Top 10 | $TCGA_{Test}$ | 0.652 | 0.570 | 0.734 | 0.662 | 0.632 | 0.570 | 0.597 | 0.310 |
| | $TCGA_{Validation}$ | 0.647 | 0.570 | 0.734 | 0.655 | 0.624 | 0.571 | 0.595 | 0.298 |
| Top 15 | $TCGA_{Test}$ | 0.674 | 0.610 | 0.738 | 0.681 | 0.651 | 0.610 | 0.628 | 0.352 |
| | $TCGA_{Validation}$ | 0.666 | 0.610 | 0.738 | 0.673 | 0.644 | 0.598 | 0.619 | 0.336 |
| Top 20 | $TCGA_{Test}$ | 0.671 | 0.603 | 0.740 | 0.679 | 0.649 | 0.603 | 0.623 | 0.348 |
| | $TCGA_{Validation}$ | 0.661 | 0.603 | 0.740 | 0.668 | 0.637 | 0.596 | 0.614 | 0.326 |
| Top 25 | $TCGA_{Test}$ | 0.677 | 0.610 | 0.745 | 0.685 | 0.657 | 0.610 | 0.630 | 0.360 |
| | $TCGA_{Validation}$ | 0.673 | 0.610 | 0.745 | 0.680 | 0.652 | 0.609 | 0.628 | 0.349 |

**Figure 13:**
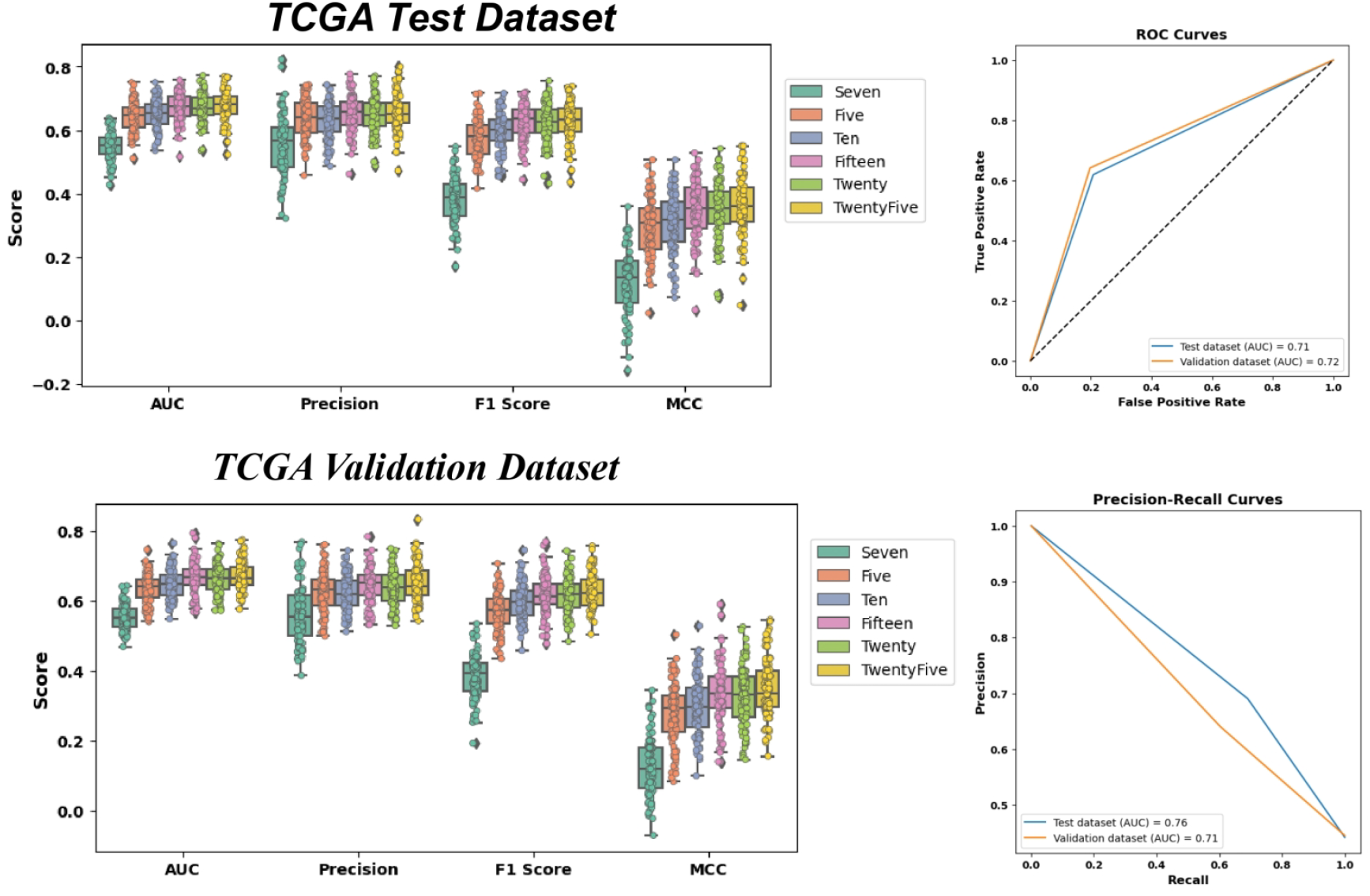
A boxplot comparison representing the performance evaluation of the expression of seven genes with the top 5, 10, 15, 20, and 25 genes based on their regulatory strength from early-stage and late-stage CRC samples. ROC and Precision-Recall curves show the predictive performance using the top 15 genes.

## 4. Discussion

Despite various advancements in the treatment of CRC, the mortality and morbidity rates continue to rise, increasing the overall disease burden [2]. This study integrates genomics information, such as CNA and mutation, with gene expression to identify the key driver genes and pathways involved in CRC progression. By integrating CNA and mutation data with gene expression, we identified a total of 3028 CGs associated with CRC, including well-known CRC driver genes such as *MYC* and *APC*. KEGG pathway analysis revealed that identified CGs were mainly involved in the endocytosis, cell cycle, spliceosome, wnt signalling pathway and mTOR signalling pathways. Endocytosis is a cellular process that involves the internalisation of extracellular substances into the cell and regulates the selective packaging of cell-surface proteins, such as receptors for cytokines and adhesion components. The endocytosis pathway has been found to be dysregulated in various cancers and acts as a key mediator between tumour cells, immune cells, and other types of cells to regulate tumour microenvironment. It plays a crucial role in the metastasis of tumours by regulating cellular processes like migration and invasion [50, 51].

The cell cycle pathway plays a crucial role in maintaining genomic integrity. Dysregulation of the cell cycle can result in uncontrolled cell proliferation and genomic instability, which are the hallmarks of cancer. Our analysis also identified the altered gene with up-regulated and down-regulated expression in CRC. The up-regulation of *CDK2, CDK4*, and E2F transcription factors family members, such as E2F1, E2F4, and E2F5, is associated with cell proliferation, which is consistent with our study. *SMAD3* and *SMAD4*, which act as tumour suppressors in various cancers, were found to be down-regulated in our analysis. Studies have also reported the role of the cell cycle in cancer metabolism and immune surveillance [52, 53, 54, 55].

The spliceosome is a complex molecule consisting of several small nuclear ribonucleoproteins. It plays an important role in splicing pre-mRNA into mature mRNA to translate into a functional protein. The alteration in the component of spliceosome can lead to the production of defects, truncated or aberrant protein products, which help in the initiation, progression, and metastasis of cancer [56].

The Wnt signalling pathway is a key player in the development and progression of CRC by regulating various biological processes, including cell proliferation, apoptosis, metabolism, inflammation, and epithelial-mesenchymal transition. Among the identified CGs, 38 genes were enriched in the Wnt signalling pathway, including 22 upregulated and 16 down-regulated. The Adenomatous Polyposis Coli (*APC*) gene and Glycogen Synthase Kinase 3 Beta (*GSK-3β*) are crucial genes of Wnt signalling pathways and act as a tumour suppressor and activator, respectively, in CRC development [57]. In our study, *APC* and *GSK-3β* genes were found to be down-regulated and up-regulated in our study. The mTOR (Mammalian target of rapamycin) signalling pathway plays an important role in the development of various cancers, including CRC. Studies have reported the involvement of the mTOR signalling pathway in signalling pathways, including the phosphoinositide-3kinase (PI3K)/AKT, apoptosis, autophagy, and metabolism [58]. We identified four genes, *CASP2, HCN4, LRRC69* and *SRD5A1*, as key driver genes associated with the progression of CRC. These genes were able to distinguish between lowrisk and high-risk groups of CRC patients. Furthermore, these genes showed a potential significance in predicting the 1-year, 3-year, and 5-year survival outcomes for CRC patients. *CASP2* (caspase-2) is a member of the caspase family and plays a crucial role in regulating various cellular processes, including genomic stability, DNA damage, activation of death receptors, and metabolic stress. In our study, *CASP2* was up-regulated with amplification as the dominant CNA type. It acts as a tumour suppressor gene by eliminating the mutated or damaged cells to reduce tumour growth. However, the function of *CASP2* is contextdependent as it was found to be up-regulated in the Human Atherosclerotic Plaques [59, 60, 61]. *HCN4*, a member of the Hyperpolarisation-activated cyclic nucleotide-gated (HCN) channel, is involved in ion transport and predominantly present in the heart and central nervous system. Studies have reported its association with neurological and cardiac-related diseases [62, 63]. Phan et al. (2017) conducted a meta-analysis of mRNA expression levels of the HCN genes family, and they observed an up-regulation of *HCN4* in leukaemia, sarcoma, lung, kidney, ovarian cancer, and thyroid cancer, whereas it was down-regulated in breast cancer [62]. Ili et al. (2020) reported the association of the *HCN4* gene with the progression and metastasis in CRC, consistent with our study [64].

*LRRC69* (leucine-rich repeat containing 69) is a leucinerich repeat (LRR) gene family member. The *LRRC69* gene in fusion with AC011997.1 has been reported to be associated with the high recurrence of breast and lung squamous cell carcinoma. AC011997.1-LRRC69 members of the LRR gene family, such as LRRC28, were reported to be dysregulated in CRC, but we did not find any study related to LRRC69 function in any type of cancer [65, 66]. SRD5A1 (Steroid 5*α*-Reductase Type I) is an enzyme that plays a key role in regulating steroid metabolism and sex hormone levels. The over-expression of the *SRD5A1* gene is reported in various cancers, including breast cancer, prostate cancer, and non-small cell lung cancer [67]. Wei et al. citewei2020steroid demonstrated in their study that the overexpression of *SRD5A1* is associated with a poor prognosis of CRC, which aligns with our study. *SRD5A1* gene was found to be connected with the identified hub genes *TEAD4* and *ETV4* through the *MYC* gene in a functional interaction network. Furthermore, some TFs are common among different biological pathways, such as *MYC* and *ETV4*, highlighting their significant role in the progression of CRC.

The WGCNA analysis identified seven hub genes associated with early and late-stage CRC: *DSCC1, ETV4, KIAA1549, NOP56, RRS1, TEAD4* and *ANKRD13B*. The diagnostic analysis, using machine learning, showed that these hub genes significantly differentiate CRC samples from normal tissues, suggesting their potential as diagnostic biomarkers for CRC. *DSCC1* (DNA replication and sister chromatid cohesion 1) plays an important role in DNA replication, spindle checkpoint and repair. The overexpression of the *DSCC1* gene has been reported in various cancers such as gastric cancer and breast cancer [68, 69, 70, 71]. In our analysis, *DSCC1* was found with amplification copy number alteration and up-regulated expression in both the altered and CRC groups compared to the normal groups. Our findings are supported by the studies of Yamaguchi et al. (2014) and Kim et al. (2019), which highlighted the role of up-regulated *DSCC1* in the progression of CRC as well as in chemoresistance [69, 72]. Their studies also highlighted the crucial role of *DSCC1* in forming a complex with CTF18-RFC (Chromosome Transmission Fidelity Factor 18-replication factor C), which is essential for DNA replication and repair. In our analysis, *RFC2* and *RFC3*, members of the RFC family, were also identified as potential CGs with up-regulated expression in CRC and showed a significant positive correlation with *DSCC1* (*r* = 0.49 and *r* = 0.61, respectively, p-value<0.05) in the CRC state. Gene regulatory network analysis revealed that the *DSCC1* gene is regulated by 127 TFs, including the E2F transcription family and hsa-miR-193b-3p miRNA. The E2F family, consisting of eight members, was found to regulate the activity of *DSCC1*, as demonstrated by Yamaguchi et al. [69]. In our analysis, we observed that E2F1, E2F4, E2F5 and E2F6 were up-regulated in CRC with amplification as the dominant copy number alteration (CNA), while E2F2 was downregulated with deletion as the dominant CNA. Among these five E2F TFs, E2F1 was found to be positively correlated (*r* = 0.57, p-value< 0.05) and E2F5 with (*r* = 0.45, p-value< 0.05) with *DSCC1* in CRC. These results highlight the significance of *DSCC1* in CRC and suggest it could be explored as a potential therapeutic target for CRC.

The ETV4, also known as ETS variant 4, is a transcription factor that belongs to the ETS transcription factor family and is involved in the regulation of various cellular processes such as cell proliferation, migration, invasion, and differentiation. Studies have shown that over-expression of ETV4 is associated with the progression of different types of cancers, including CRC [73, 74].

KIAA1549-BRAF fusion in pilocytic astrocytoma, breast carcinoma, and sarcomas. In our study, both the *KIAA1549* and *BRAF* were found to be up-regulated in the altered CRC state as compared to the normal state. In CRC, *KIAA1549* has been found to be highly expressed and has been shown to promote the proliferation and migration of CRC cells [75].

*NOP56* is an essential and highly conserved nucleolar protein involved in ribosome production and plays an important role in processing rRNA precursors and promoting the synthesis of mature rRNA. *NOP56* act as a central point for various signalling pathways such as NF-*κ*B signalling, JAK/STAT signalling and mTOR signalling pathway. Studies have shown that *NOP56* expression is significantly up-regulated in certain cancers such as acute myeloid leukaemia, diffuse large B-cell lymphoma, non-small cell lung cancer, pancreatic cancer, colorectal cancer, breast cancer, prostate cancer, and acute lymphoblastic leukaemia [76].

## 5. Conclusion

This multi-omics study integrated copy number alterations (CNA), mutations, and gene expression data, resulting in the identification of 3,028 candidate genes associated colorectal cancer (CRC). Among these, several known CRC driver genes such as *MYC* and *APC* were highlighted. A further pathway analysis indicated that these candidate genes are signalling pathways, including endocytosis, cell cycle regulation, Wnt signalling, and mTOR signalling pathways. Furthermore, our extended study on these candidate genes revealed seven hub genes, *DSCC1, ETV4, KIAA1549, NOP56, RRS1, TEAD4* and *ANKRD13B*, were identified through the Weighted Gene Co-expression Network Analysis (WGCNA). These hub genes demonstrated significant potential as diagnostic biomarkers for CRC when used for building diagnostic models using machine learning. Survival analysis pinpointed four genes, *CASP2, HCN4, LRRC69* and *SRD5A1*, as potential key driver genes associated with the progression of CRC and can be considered as prognostic biomarkers.

This multi-omics and machine learning based comprehensive identification of key driver genes, enriched pathways, and pivotal hub genes not only enhance understanding of the molecular mechanisms involved in CRC progression but also holds promise for the development of more effective diagnostic and prognostic tools. This study provides substantial support for the study of a complex and life-threatening disease like CRC. While there is still a long journey to cover in diagnosis and treatment of CRC, this study could certainly aid in the community’s thinking towards developing personalised therapeutics.

## Supplementary Files

We provide additional supporting results in the supplementary document.

## Data and code availability

The datasets used in our study are available in the public databases, as described in the manuscript. Nevertheless, we release some processed datasets publicly for convenience and reproducibility. These processed datasets and codes used in our experiments are available at: https://github.com/Mohita1304/multi-omics-Analysis.

## Acknowledgements

The authors sincerely thank the Birla Institute of Technology and Science, Pilani, K.K. Birla Goa Campus, for supporting this work.

## CRediT authorship contribution statement

**Mohita Mahajan**: Writing – original draft, Writing – review & editing, Methodology, Data curation, Software, Formal analysis. **Subodh Dhabalia**: Software, Data curation. **Tirtharaj Dash**: Writing – review & editing, Supervision. **Angshuman Sarkar**: Writing – review & editing, Supervision. **Sukanta Mondal**: Conceptualization, Writing – review & editing, Supervision, Investigation.

