## Supplementary Files for "A comprehensive multi-omics study reveals potential prognostic and diagnostic biomarkers for colorectal cancer"

- Risk $Score_{GSE29623}$ = (-0.546)*CASP2 + (-0.499)*HCN4 + (0.944)*LRRC69+ (0.590)*SRD5A1
- Risk $Score_{GSE17536}$ = (-0.111)*CASP2 + (-2.178)*HCN4 + (0.421)*LRRC69+ (0.310)*SRD5A1

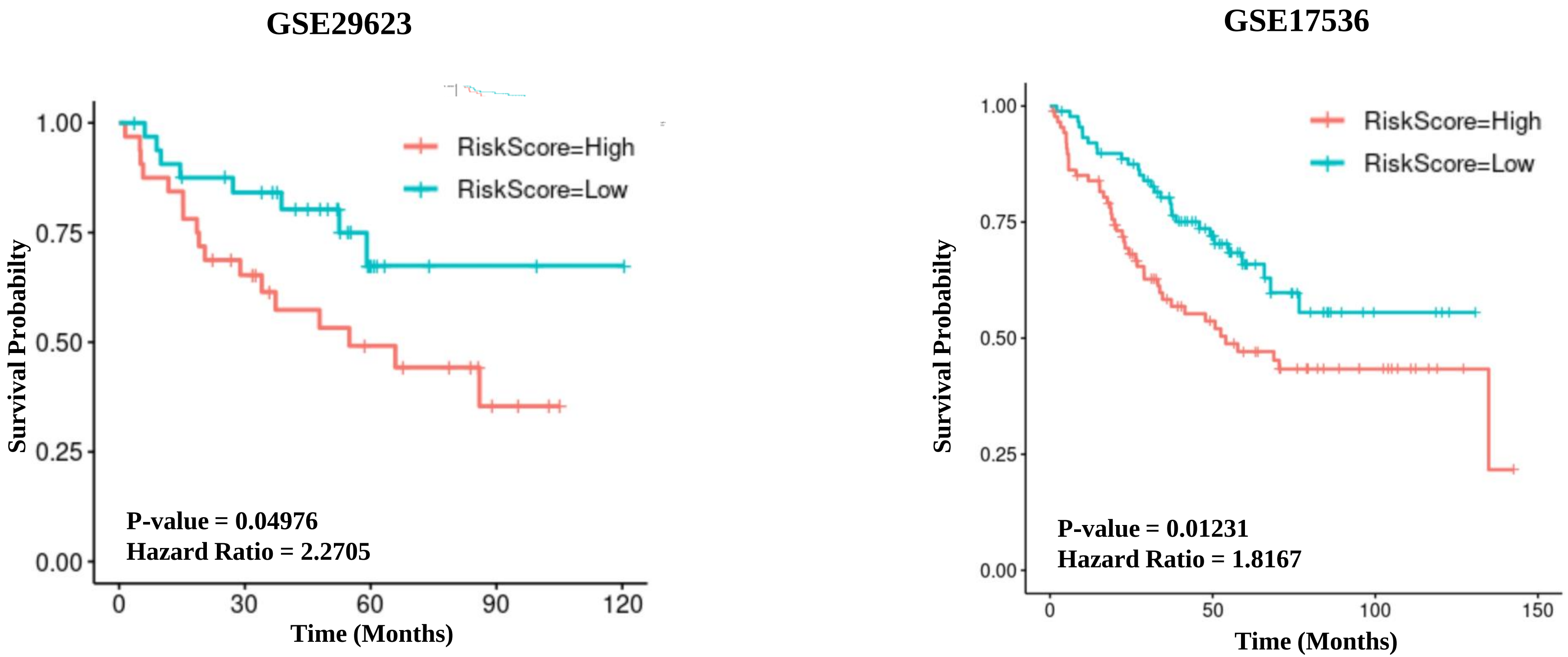


**Figure S1:** Survival curve for CRC patients with low-risk (RiskScore < median) and high-risk (RiskScore > median) groups.

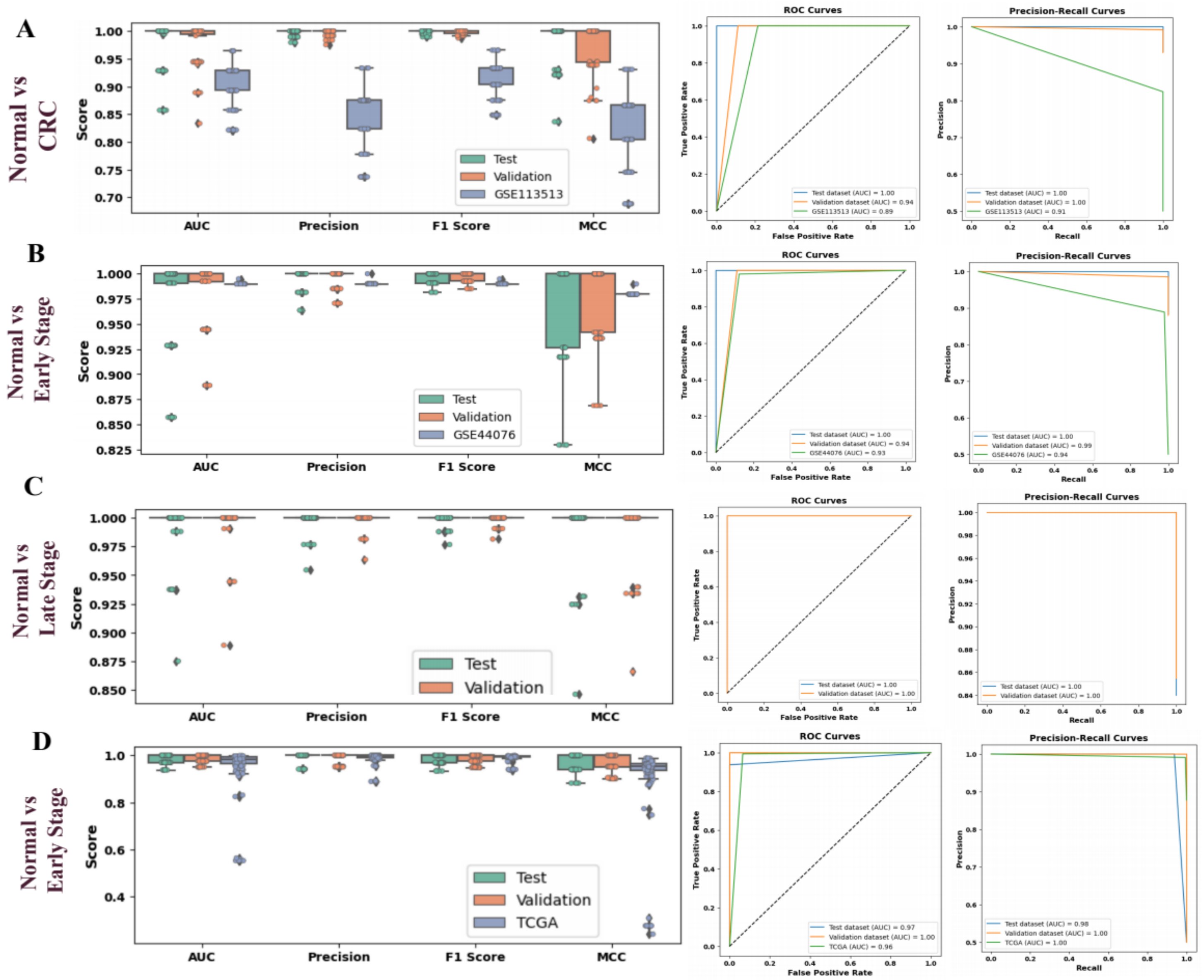


**Figure S2.** Diagnostic performance of the seven hub genes in discriminating the normal sample to the early and late stage and CRC samples using the Random Forest model. (A-C) TCGA datasets used for training the prediction model (D) GSE44076 dataset was used to train the model for normal vs. Early stage CRC.

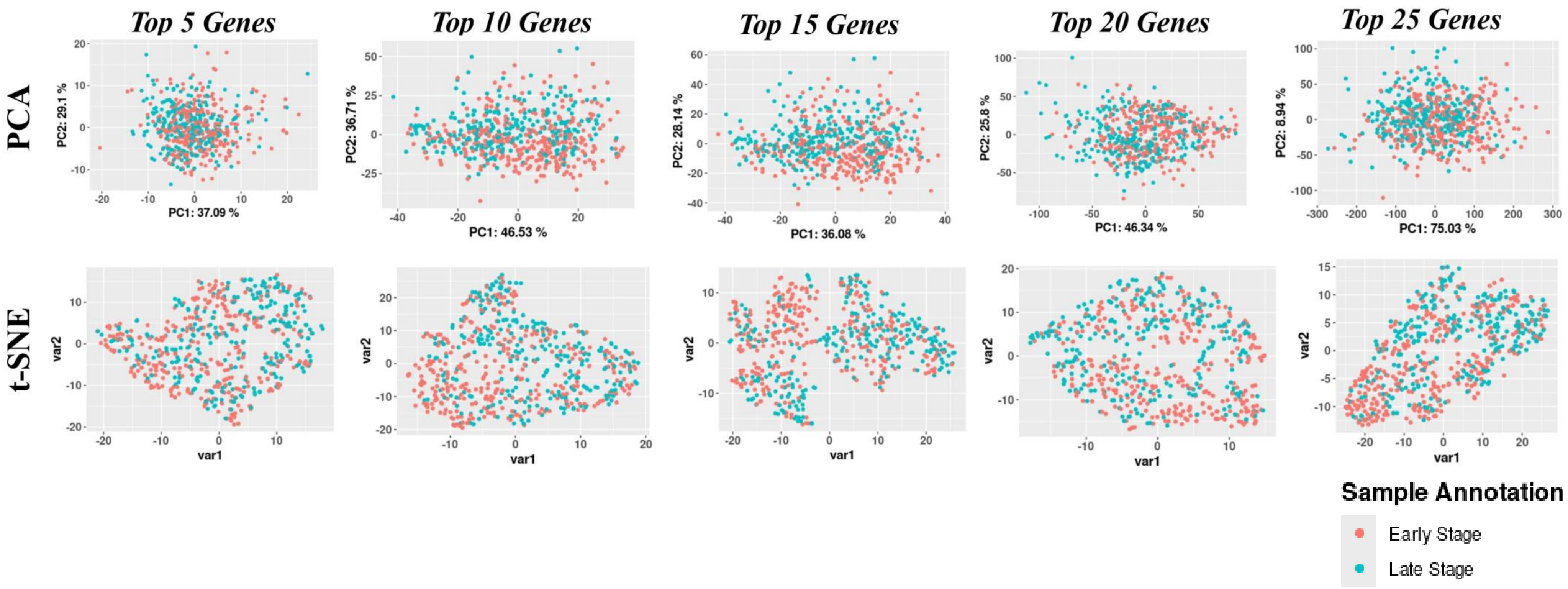


**Figure S3.** PCA and t-SNE analysis representing the discrimination ability of the selected top ranked genes by using mRMR algorithm based on the regulatory strength of the genes.

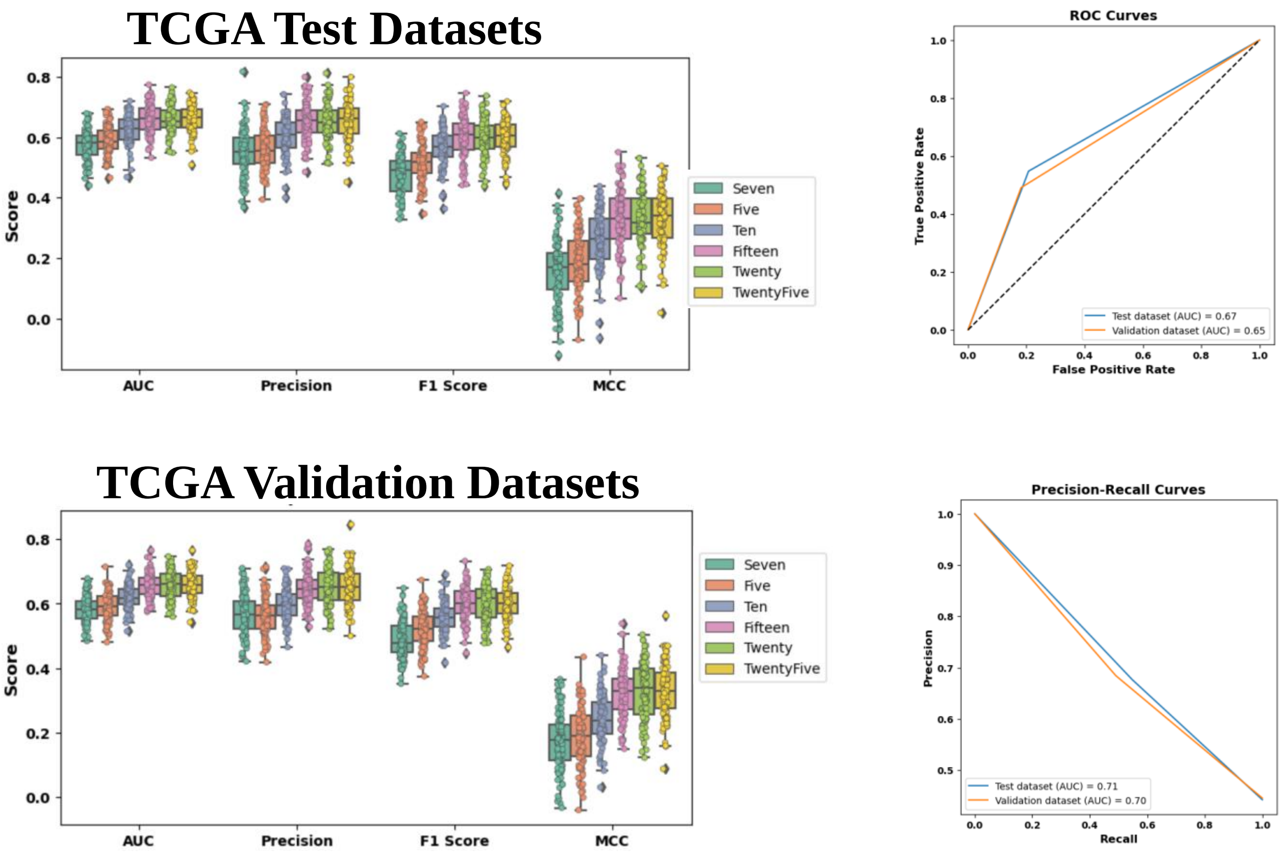


**Figure S4.** A boxplot comparison representing the performance evaluation (Random Forest) of the expression of seven genes with the top ranked 5, 10, 15, 20, and 25 genes based on their regulatory strength from early-stage and late-stage CRC samples. ROC and Precision-Recall curves showing the predictive performance using the top 15 genes.

**Table S1**. Cross-referencing results of the identified CGs with established CRC-associated genes from the DisGeNET database and oncogenic driver genes and driver mutation from the OncoVar database.

| Association Type | Database | CGs |
|---|---|---|
| Disease-gene | DisGeNET | ACAP1, SMAD3, CDC14A, CCND2, MYC, BUB1, SMAD4, SNRPB2, BCAS2, SRSF6, SNRNP200, TCF7L2, RNF43, NFATC1, AXIN2, SFRP4, APC, RHEB, FLCN, SESN2, ATP6V1E1, BRAF, AURKA, SLC11A2, CAD, PMM2, NME2, GGH, UMPS |
| Onco driver gene | OncoVar | MSH2, MSH6, PCBP1, PMS1, PMS2, SFRP4, BRAF, EIF3E, AXIN1, RNF43, AXIN2, SALL4, CD58, PIK3R1, APC, TCF7L2, MAX, B2M, SMAD3, MAP2K4, SMAD4 |
| Onco driver mutation | OncoVar | KIF14, LBR, KIF26B, PREB, XPO1, RPIA, MAP4K4, ERCC3, CERS6, SF3B1, ITGB5, TRIO, ADAMTS12, KATNA1, TULP4, MAD1L1, WIPI2, PMS2, GRB10, AUTS2, LIMK1, NPTX2, POT1, GRM8, BRAF, TAS2R38, EZH2, SLC4A2, PAXIP1, SHH, RNF170, CHD7, STK3, FZD6, SLC25A32, PTK2, TONSL, C9orf72, PRMT3, PEX5, XPOT, SLC7A1, BRCA2, ATP7B, DIS3, F7, ABCC1, CDR2, TUFM, SRCAP, VAC14, WDR59, SPG7, ACACA, CDK12, EFTUD2, DOCK6, KMT2B, SHKBP1, SLC1A5, PRPF31, U2AF2, STK35, LPIN3, TTPAL, ZMYND8, OGT, MCTS1, GABRE, CAMTA1, KIF1B, MTFR1L, AGL, RAF1, RHOA, FLNB, LIAS, SLC30A9, SLC4A4, G3BP2, SH3RF1, APC, CTNNA1, PPP3CC, NUGGC, GSR, VCL, TCTN3, WBP1L, TCF7L2, RAB1B, DAAM1, MAX, PSEN1, ALDH6A1, SPTLC2, RPS6KA5, TRAF3, CDC42BPB, MAPK6, TLN2, HCN4, ARNT2, INPP5K, MAP2K4, TUBB6, KCTD1, IER3IP1, SMAD4, FECH, ATP8B1, NEDD4L, SYNJ1, ADARB1, MTMR3, PPARA |

**Table S2**. Average Performance Metrics of the Random forest Model for Predicting CRC Status.

| Study | Validation | AUC | Sensitivity | Specificity | Accuracy | Precision | Recall | F1 Score | MCC |
|---|---|---|---|---|---|---|---|---|---|
| Normal vs. CRC | $TCGA_{Test}$ | 0.985 | 1.000 | 0.97 | 0.998 | 0.998 | 1.000 | 0.999 | 0.981 |
| | $TCGA_{Validation}$ | 0.984 | 1.000 | 0.967 | 0.997 | 0.998 | 0.999 | 0.998 | 0.978 |
| | GSE113513 | 0.904 | 1.000 | 0.857 | 0.904 | 0.843 | 1.000 | 0.914 | 0.825 |
| Normal vs. Early Stage | $TCGA_{Test}$ | 0.949 | 1.000 | 1.000 | 0.983 | 0.988 | 0.993 | 0.990 | 0.907 |
| | $TCGA_{Validation}$ | 0.986 | 0.998 | 0.963 | 0.996 | 0.996 | 0.999 | 0.998 | 0.980 |
| | GSE44076 | 0.990 | 1.000 | 1.000 | 0.990 | 0.990 | 0.990 | 0.990 | 0.980 |
| | $GSE44076_{Test}$ | 0.984 | 0.973 | 0.996 | 0.984 | 0.996 | 0.973 | 0.984 | 0.97 |
| | $GSE44076_{Validation}$ | 0.985 | 1.000 | 1.000 | 0.985 | 0.992 | 0.978 | 0.984 | 0.970 |
| | TCGA | 0.949 | 1.000 | 1.000 | o.983 | 0.988 | 0.993 | 0.990 | 0.907 |
| Normal vs. Late-Stage | $TCGA_{Test}$ | 0.996 | 0.999 | 0.993 | 0.998 | 0.999 | 0.999 | 0.999 | 0.993 |
| | $TCGA_{Validation}$ | 0.996 | 1.000 | 0.991 | 0.999 | 0.999 | 1.000 | 0.999 | 0.995 |
